# Narrowband gamma oscillations propagate and synchronize throughout the mouse thalamocortical visual system

**DOI:** 10.1101/2022.05.19.491028

**Authors:** Donghoon Shin, Kayla Peelman, Joseph Del Rosario, Bilal Haider

## Abstract

Rhythmic oscillations of neural activity permeate sensory systems. Studies in the visual system propose that broadband gamma oscillations (30 – 80 Hz) facilitate neuronal communication underlying visual perception. However, broadband gamma oscillations within and across visual areas show widely varying frequency and phase, providing constraints for synchronizing spike timing. Here, we analyzed data from the Allen Brain Observatory and performed new experiments that show narrowband gamma (NBG) oscillations (50 – 70 Hz) propagate and synchronize throughout the awake mouse thalamocortical visual system. Lateral geniculate (LGN) neurons fired with millisecond precision relative to NBG phase in primary visual cortex (V1) and multiple higher visual areas (HVAs). NBG in HVAs depended upon retinotopically aligned V1 activity, and neurons that fired at NBG frequencies showed enhanced functional connectivity within and across visual areas. Remarkably, LGN ON versus OFF neurons showed distinct and reliable spike timing relative to NBG oscillation phase across LGN, V1, and HVAs. Taken together, NBG oscillations may serve as a novel substrate for precise coordination of spike timing in functionally distinct subnetworks of neurons spanning multiple brain areas during awake vision.

## Introduction

Oscillations of neural activity are thought to play an important role in both representing and communicating sensory information across the brain. Extensive studies in visual cortex have shown that broadband gamma (30 – 80 Hz) oscillations vary with visual stimulus features and visual task performance (Fries et al., 2001; Gray et al., 1989), leading to the hypothesis that synchronized gamma activity across visual brain areas facilitates neuronal communication underlying perception (Fries, 2015; Singer and Gray, 1995). However, recent work identifies several limitations for broadband gamma oscillations to both represent sensory stimuli and synchronize communication across brain areas (Kohn et al., 2020; Ray and Maunsell, 2015). First, broadband gamma oscillations measured simultaneously across brain areas show high variability in frequency, amplitude, and phase. Second, there is no “central clock” that coordinates broadband gamma oscillations across brain areas at millisecond timescales. Third, broadband gamma oscillations typically emerge slowly after stimulus onset and then fluctuate continuously with stimulus features. All these factors pose constraints for maintenance of spike timing precision and synchronization across widespread visual areas. Instead, broadband gamma oscillations may primarily reflect timescales of local cortical excitation and inhibition (Buzsaki and Wang, 2012; Cardin, 2016; Sohal, 2016). If, however, an oscillation acts primarily to coordinate visual activity across brain areas, then it should 1) show consistent frequency and phase across regions, 2) show synchronization at the visual input layers across these regions, and 3) enforce neurons to fire at distinct oscillation phases according to visual feature preferences.

Recent studies have unveiled a novel narrowband gamma (NBG) oscillation in the mouse visual system. Unlike broadband gamma activity, NBG in primary visual cortex (V1) shows a highly stereotyped oscillation frequency (central peak between 50 – 70 Hz), narrow bandwidth (5 – 7 Hz), and is not generated by visual stimulus features. NBG in mouse V1 arises spontaneously during wakefulness (Niell and Stryker, 2010), disappears in total darkness, and propagates directly from lateral geniculate nucleus (LGN; Saleem et al., 2017) and likely from retina (Storchi et al., 2017), echoing findings from the cat visual system (Castelo-Branco et al., 1998; Neuenschwander and Singer, 1996). Remarkably, NBG in mouse V1 also varies with arousal and behavioral state (Haider et al., 2013; Niell and Stryker, 2010; Saleem et al., 2017; Vinck et al., 2015), and its strength predicts visual perceptual performance (Speed et al., 2019). This suggests that NBG could serve to coordinate activity across visual areas underlying perception. However, it is unknown if NBG activity propagates beyond V1 to higher visual areas (HVAs) that are also involved in visual perception. If NBG does invade HVAs, it is unknown how this depends on V1 activity, if NBG is tightly synchronized across visual cortical areas, or if NBG enforces spike timing of neurons depending on their visual stimulus preferences. Answering these questions requires large-scale, simultaneous neural recordings from LGN, V1, and HVAs, perturbations of NBG across areas, and a framework for detecting and quantifying population and single-neuron level NBG activity.

Here we addressed these questions by analyzing the Allen Brain Observatory Visual Coding dataset of multi-site simultaneous Neuropixels recordings (Allen Brain Observatory, 2019; Siegle *et al*., 2021), and by performing recordings from HVAs during simultaneous optogenetic inactivation of V1. We found strong evidence for NBG activity propagation and synchronization throughout the mouse thalamocortical visual system. We found many neurons in LGN, V1, and HVAs that showed significantly synchronized NBG spiking, and these neurons showed greater likelihood for pairwise functional interactions than neurons lacking NBG activity. NBG in LGN was tightly synchronized with local field potential (LFP) oscillations in the input layers of V1 and HVAs, and NBG in HVAs showed retinotopic dependence on V1 activity. Surprisingly, LGN neurons preferring dark (OFF) versus bright (ON) visual stimuli fired spontaneously at distinct phases of NBG oscillations; these feature and phase preferences in LGN spikes were aligned with NBG activity in V1 and multiple HVAs according to their position along the anatomical hierarchy. Together, these findings show that NBG oscillations effectively synchronize spiking in functionally distinct groups of neurons throughout the awake mouse visual system, identifying a novel potential substrate for rapid communication and coordination of visual information across the brain.

## Results

### Correlated NBG spiking across LGN, V1, and HVAs

We first verified NBG communication from LGN to V1 in the Allen Brain Observatory – Visual Coding-Neuropixels dataset, and then found evidence for correlated NBG spiking across LGN, V1, and HVAs. Our starting point was to examine simultaneous Neuropixels recordings of spikes in LGN and V1 (Fig. 1A). Consistent with prior reports, (Saleem *et al*., 2017; Schneider et al., 2021), many individual LGN neurons showed NBG power (between 50 – 70 Hz) in their spike autocorrelograms (ACGs; Fig. S1). Further, we found that cross-correlograms (CCGs) of many LGN-V1 pairs also showed correlated spiking oscillating at NBG frequencies (Fig. 1B). We identified NBG neurons as those with CCGs that fulfilled several quantitative metrics of significant NBG power (see Methods and Fig. S1). Correlated NBG spiking between LGN and V1 neuron pairs was highly specific: the same LGN neuron could show significant NBG firing synchronized with some V1 neurons (Fig. 1B), but not others recorded simultaneously and just tens of microns away on the probe (Fig. 1C). In this example session, we found many significant NBG CCGs between LGN - LGN neuron pairs (Fig. 1D; n = 57), and between LGN - V1 neuron pairs (n = 21), and LGN - HVA neuron pairs (n = 8). CCGs between these same NBG neurons and other neurons recorded simultaneously showed no significant NBG power in any CCG (Fig. 1E), ruling out global synchronization and instead suggesting that only specific neurons show correlated NBG firing across the visual system. Across all sessions with LGN single unit recordings in the Allen dataset (n = 32 recordings), 35% of all LGN neurons were classified as NBG neurons (n = 455; Table S1), while the remainder were classified as non-NBG neurons.

**Figure 1.**
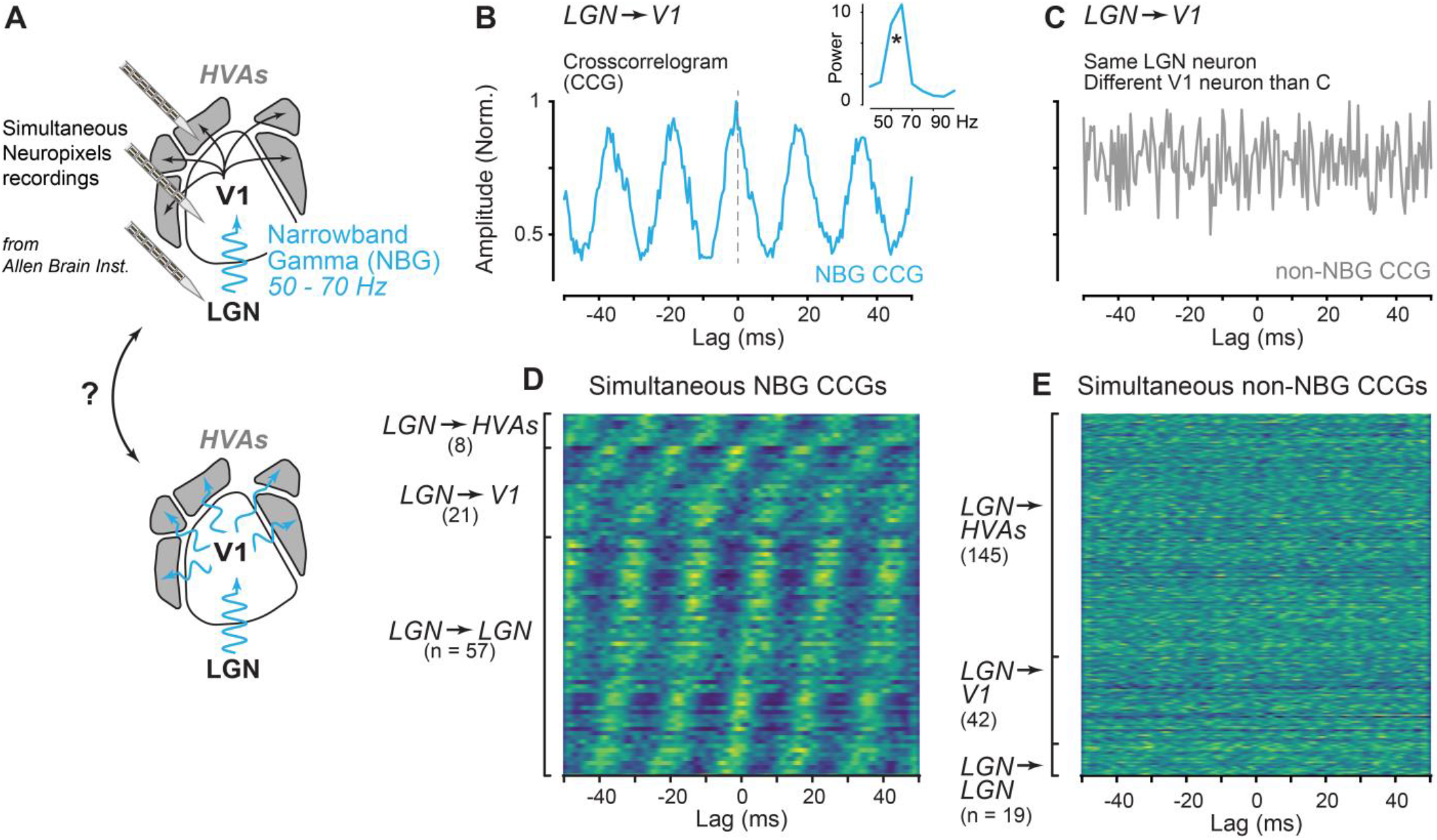
Synchronized narrowband gamma (NBG) activity across thalamocortical visual areas. **A**. Simultaneous multisite Neuropixels recordings from lateral geniculate nucleus (LGN), primary visual cortex (V1), and higher visual areas (HVAs) from the Allen Brain Observatory Visual Coding - Neuropixels dataset were examined for evidence of NBG oscillations from LGN to V1 (top), and for evidence of NBG activity across LGN, V1, and HVAs (bottom). **B**. Example spike cross-correlogram (CCG) between a NBG neuron in LGN and a V1 neuron during spontaneous activity in an example session (170 mins of simultaneous recording with ∼161k LGN spikes and ∼393k V1 spikes). Neurons in each area were classified as NBG neurons if their autocorrelograms showed significant spectral power in the 50 – 70 Hz range, or if the CCG showed significant NBG power (inset; see Methods and Fig. S1). **C**. Same NBG LGN neuron in B, and CCG with a non-NBG neuron in V1 during spontaneous activity during the same session as B. Neurons without significant 50-70 Hz power in autocorrelations or cross-correlations classified as non-NBG. Non-NBG neuron recorded on probe site adjacent to the one in B. **D**. Cross-correlograms between NBG LGN neuron in B, and all other simultaneously recorded NBG neurons in LGN (n = 57), V1 (n = 21) and higher visual areas (n = 8) in same example session. Heatmap shows each CCG normalized to peak amplitude (yellow) and sorted by peak lag for visualization of oscillatory firing. Plot excludes one NBG cell in LGN for display. **E**. Cross-correlograms between the same NBG LGN neuron as B-D and all simultaneously recorded non-NBG neurons. Same conventions and recording session as D.

We performed several additional control measures to confirm that identification of NBG LGN neurons by CCGs depended upon spike timing. First, jittering spike times by ±20ms significantly reduced NBG power in CCGs, and in no case did jittered CCGs lead to misidentification of NBG neurons according to our criteria (<0.001% false positive rate in n = 30 pairs from 15 recording sessions with >10k jittered CCGs per pair; see Methods and Fig. S1). Second, computing spike CCGs between NBG LGN neurons from one recording session (n = 34) and all neurons from a different recording (n = 242 neurons) never yielded a false positive classification of NBG neuron pairs, despite the LGN neurons showing significant NBG power in CCGs within session (>8k pairwise CCGs). Third, identification of NBG neurons using CCGs showed high overlap with identification based solely on spike autocorrelograms (ACGs; Fig. S1), and our main results were unchanged even when we restricted analysis solely to NBG neurons identified with ACGs, as will be evident later.

NBG local field potential (LFP) oscillations across V1 and HVAs were synchronized with spikes of NBG LGN neurons. We examined V1 LFPs triggered on simultaneously recorded spikes of NBG neurons in LGN (Fig. 2A). We used current source density analysis (Fig. S2) to localize the earliest current sink that corresponds to input layer 4 (L4) of V1, then calculated the spike triggered LFP (stLFP) at this site relative to NBG LGN neuron spikes during spontaneous activity. An example V1 L4 stLFP showed clear and significant NBG oscillations (Fig. 2CB; 55.6 Hz), with LGN spikes preceding the LFP trough by 5.6 ms. We then identified the putative functional input layers of HVAs in the same way (Fig. S2) and calculated the stLFP in HVAs relative to NBG LGN spikes. As expected, the stLFP power was greatest in V1 (Fig. 2C) but also showed elevated power in the lateral HVAs (RL, LM) compared to the medial ones (PM, AM; schematic in Fig. 2A). Importantly, stLFP NBG power was significantly reduced when triggered from non-NBG LGN neuron spikes in the same recordings (−6.9 ± 6.9 dB reduction on average), a significant difference in all areas except PM (V1: *p* < *1e-3*; RL: *p* < *0*.*04*; LM: *p*< *3e-4*; AL: *p* < *6e-4*; AM: *p = 0*.*03;* PM: *p = 1;* Wilcoxon rank sum tests). We then plotted stLFP power versus hierarchy scores that directly reflect feedforward versus feedback anatomical connectivity (Harris et al., 2019), and found a significant correlation between the anatomical hierarchy and stLFP NBG power (Fig. 2D; r = -0.9, *p = 0*.*01*). Further, this steep dependence of stLFP NBG power on the anatomical hierarchy (linear slope of -23.8 dB; Fig. 2D) was significantly weaker when triggered on non-NBG LGN neurons (linear slope of -11.0 dB, *p* < *4e-5*).

**Figure 2.**
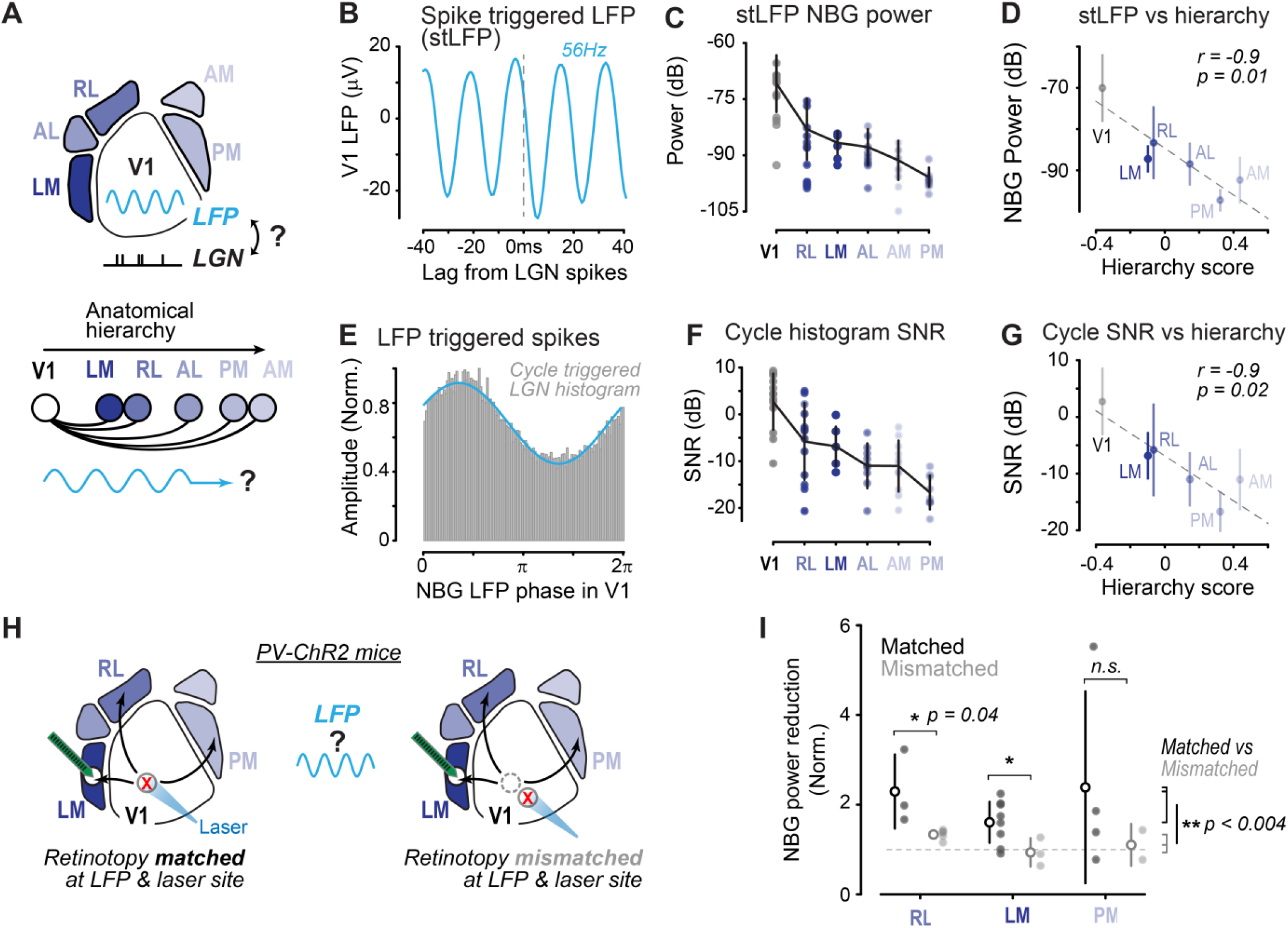
Synchronized NBG across the hierarchy of HVAs depends upon retinotopic V1 activity. **A**. *Top*, schematic of measuring V1 LFP triggered on LGN spikes. *Bottom*, anatomical hierarchy of V1 and HVAs. Hierarchical order indicated by color saturation (LM, lateromedial area; RL, rostrolateral area; AL, anterolateral area; PM, posteromedial area; AM, anteromedial area), based on (Siegle et al., 2021). **B**. Example spike triggered LFP (stLFP) in V1 layer 4 (L4) triggered by simultaneously recorded NBG LGN neuron spikes. Peak LFP power at 55.6Hz. V1 L4 identified with current source density analysis (Fig. S2). **C**. stLFP NBG power across V1 and HVAs aligned to NBG LGN neuron spikes (n = 400 neurons, 14 sessions) sorted from highest to lowest stLFP power (mean ± SD), circles show individual session means. **D**. Average stLFP NBG power of V1 and HVAs (mean ± SD) sorted by anatomical hierarchy scores (see Methods). Significant correlation with anatomical hierarchy (Pearson rho = -0.90; *p = 0*.*0142*). **E**. Example NBG LGN neuron spike histogram aligned to V1 NBG LFP cycle phase. Blue curve shows fitted cosine used to estimate cycle histogram signal to noise ratio (SNR, see Methods). **F**. Cycle histogram SNR of V1 and HVAs across all recording sessions. Same conventions as C. **G**. Significant correlation of cycle histogram SNR with anatomical hierarchy scores. (Pearson rho = -0.89; *p = 0*.*0162*). Same conventions as D. **H**. HVA recordings with optogenetic inactivation of V1, when retinotopy is matched (*left)*, or mismatched (*right*) at stimulation and recording sites. V1 inactivation via channelrhodopsin (ChR2) in parvalbumin (PV) inhibitory neurons. See Fig. S2 for V1 and HVA targeting via intrinsic signal imaging maps. **I**. Significantly greater NBG power reduction across HVAs during retinotopically matched V1 inactivation (2.0 ± 1.0; all areas) versus mismatched inactivation (1.2 ± 0.3; *p = 0*.*0034*, Wilcoxon rank sum; n = 26 recordings plus inactivation in 5 mice). Significantly greater NBG reduction for matched V1 inactivation in RL and LM (*p = 0*.*04* for both, Wilcoxon rank sum). NBG power reduction normalized by broadband gamma reduction (see Methods).

Is NBG phase in V1 and HVAs coupled to periodicity of LGN spike trains? We filtered the LFP in V1 and HVAs between 50 – 70 Hz, and then computed histograms of NBG LGN neuron spikes aligned to peaks of cortical LFP filtered for NBG frequencies (Methods). We found clear periodicity in LGN spiking relative to continuous NBG LFP phase in V1 (Fig. 2E; example session cycle histogram). We quantified the amount of sinusoidal power in cycle histograms by computing their signal to noise ratio (SNR; see Methods) and found that LGN spike cycle histograms showed greatest SNR in V1, followed by lateral (RL, LM) then medial HVAs (PM, AM; Fig. 2F). We again used non-NBG LGN neurons as an internal control, and found cycle histograms of NBG neurons showed significantly greater power relative to cortical LFP in all areas (V1: *p* < *3e-5*; RL: *p* < *2e-3*; LM: *p*< *2e-4*; AL: *p* < *4e-3*; AM *p* < *4e-3*) except PM (p = 0.14), and significantly steeper relationship of histogram SNR power in NBG neurons relative to non-NBG neurons in the same recordings (−19.9 dB across areas for NBG neurons vs -4.6 dB for non-NBG neurons, *p* < *5e-7*). We plotted cycle histogram SNR power versus anatomical hierarchy scores, and again found a significant correlation (Fig. 2H; r = -0.9; *p = 0*.*02*). These findings with stLFP and cycle histograms suggest that NBG activity in LGN and V1, the major source of feedforward cortical input to the HVAs (Siegle *et al*., 2021), may contribute to propagation of NBG activity across the visual cortical hierarchy.

Indeed, we found that the strength of NBG across HVAs depended upon retinotopically aligned feedforward input from V1. We performed new experiments (not part of the Allen Brain Observatory dataset) to verify NBG in LGN, and causally assess contributions of V1 to NBG activity in HVAs. We first confirmed strong and prevalent NBG spiking in our recording conditions with Neuropixels recordings from LGN (nearly 25% of all LGN neurons classified as NBG neurons; 196 / 805 in n = 7 mice; 26 recordings; Fig. S2), supporting our previous observations of NBG in V1 (Speed et al., 2019). We then performed experiments that optogenetically inactivated retinotopically defined regions of V1 (by driving channelrhodopsin (ChR2) in parvalbumin (PV) interneurons) while simultaneously recording in downstream HVAs RL, LM and PM (n = 5 mice;26 experiments; see Fig. S2 for V1 and HVA visualization and targeting). Inactivation of V1 significantly reduced NBG in HVAs. Across all experiments, residual NBG LFP power was reduced more than 3-fold (3.3 ± 1.9) versus interleaved control trials; in contrast, overall broadband gamma LFP power (30 – 90 Hz; excluding NBG at 50 – 70 Hz) was reduced to a significantly lesser extent (2.0 ± 0.6; *p = 0*.*0031*, Wilcoxon rank sum; see Methods). We accounted for these concerted activity changes by normalizing the residual NBG power reduction by overall broadband gamma power reduction within each experiment (Fig. 2I). Remarkably, the normalized NBG power reduction in HVAs depended upon retinotopic alignment with V1: inactivating V1 sites with receptive fields (RFs) that matched those at the HVA recording sites (10.2 ± 2.5° apart; Methods) led to two-fold reductions of NBG power (2.0 ± 1.0), significantly greater than inactivation of V1 sites with RFs distant from the HVA recording sites (1.2 ± 0.3 reduction with RFs 47.4 ± 2.6° apart; *p = 0*.*0034*, Wilcoxon rank sum). Taken together, these results show that the strength of NBG activity propagation across multiple HVAs depends upon retinotopically aligned feedforward input from V1. These effects relied on inactivating V1 neural populations then measuring effects on NBG activity in HVA populations. We next sought evidence for functional connectivity between individual NBG neurons spanning visual areas in the Allen Brain Observatory data.

We found a significantly greater probability of pairwise functional connectivity among NBG neurons versus non-NBG neurons within and across areas. We first assessed pairwise functional connectivity between pairs of LGN and V1 neurons (Fig. 3A). We examined all pairwise CCGs and defined significant functional interactions in CCGs that displayed statistically significant peaks in the CCG that were delayed by 1 – 4.5ms (Methods; Table S1), as in prior studies (Senzai et al., 2019; Stark and Abeles, 2009). We compared functional interactions between NBG neurons to those between all non-NBG neurons (Fig. 3B-D), and also between non-NBG with high firing rates (see Methods), since these provide an upper bound for estimating functional interactions via spike cross correlations (de la Rocha et al., 2007). Across all LGN – V1 pairs, we found a significantly greater probability of functional connectivity among NBG neurons (0.60%; 26 / 4,305) compared to all non-NBG neurons (Fig. 3B; 0.009% 7 / 78,283; *p* < *1e-37*, Binomial t-test), and compared to non-NBG neurons with high firing rates (0.02% 1/5771; *p* < *1e-30*) all in the very same recording sessions (n = 58 sessions). These trends were even more pronounced within V1, where NBG neuron pairs showed a nearly 5% probability of functional connectivity, significantly greater than among all non-NBG neurons (1% probability; Fig. 3C; *p* < *2e-42)* and non-NBG neurons with high firing rates (2.7% probability; *p* < *4e-9*). We then examined pairwise interactions between LGN and HVAs (aggregated together; Table S1) and found that here too the probability of functional connectivity was significantly greater among NBG versus all non-NBG neurons (Fig. 3D; 1% versus 0.005%, *p* < *1e-33*) and versus non-NBG neurons with high firing rates (0.01%, *p* < *1e-28*). Expanding our view of CCGs to include statistically significant peaks at zero lag (suggesting common input to both neurons; see Methods), we found further evidence for preferential interactions among NBG neurons: nearly 30% of V1 NBG pairs (409 /1,332) showed evidence for significant common input, versus 5% among all non-NBG V1 pairs (6,302 / 135,219; *p*<*1e-208*), with similar trends in LGN-V1 pairs (NBG: 0.35%,15 / 4,305; non-NBG: 0.02%, 13 / 78,283; *p* < *1e-14*) and LGN-HVA pairs (NBG: 0.61%, 10 / 1641; non-NBG: 0.02%, 49 / 248,209; *p* < *1e-12*).

**Figure 3.**
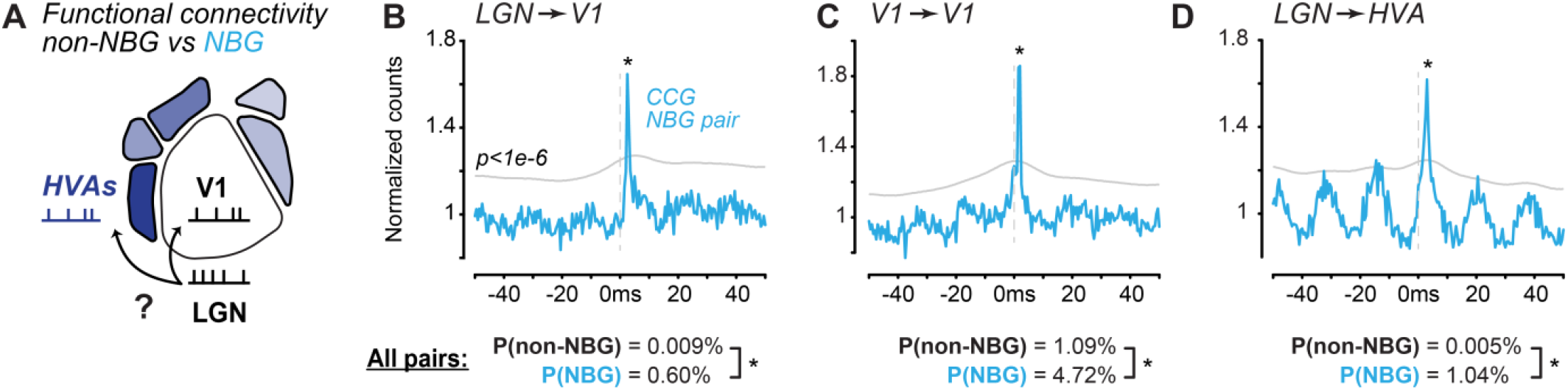
NBG neurons show higher probability of functional connectivity across visual areas. **A**. Spike trains of LGN, V1, and HVA neurons were examined for evidence of functional connectivity. **B**. *Top*, pairwise spike cross-correlogram (CCG, 0.5ms bins) of NBG neuron in LGN (∼79k spikes) and NBG neuron in V1 (∼121k spikes; 161 mins of recording). Grey line shows significance threshold for expected Poisson process (∼*p ≤ 1e-6*; see Methods). CCGs with >2 consecutive threshold crossings in 1.5–4ms classified as functional connectivity pairs. *Bottom*, across all LGN – V1 pairs, probability of functional connectivity among NBG pairs (0.60%; 26 / 4,305 pairs) was significantly greater than all non-NBG pairs (0.009%; 7 / 78,283 pairs; *p* < *1e-37*, Binomial t-test, same throughout figure). 1,309 LGN and 3,694 V1 neurons in 58 recording sessions. NBG pairs also showed significantly greater probability of common input (0.35%) versus non-NBG pairs (0.02%; *p* < *1e-14*; see Results and Methods). **C**. As in B, for pairwise correlations within V1. Probability of functional connectivity among V1 NBG pairs (4.72%; 126 / 2,664) significantly greater than all non-NBG pairs (1.09%, 2,848 / 270,438; *p* < *1e-42*). 3,694 V1 neurons in 58 recording sessions. NBG pairs also showed significantly greater probability of common input (30.70%) than all non-NBG pairs (4.66%; *p* < *1e-14*; see also Table S3). **D**. As in B, for pairwise correlations between LGN and HVAs. Example shows CCG between NBG neurons in LGN and RL. Probability of functional connectivity among LGN-HVA NBG pairs (1.04%, 17 / 1,641) significantly greater than all non-NBG pairs (0.005%, 12 / 248,209; *p* < *1e-33*). 1,309 LGN and 12,435 HVA neurons in 58 sessions. NBG pairs also showed significantly greater probability of common input (0.61%) versus all non-NBG pairs (0.02%; *p* < *1e-11*; Table S3). All data in B – D from Allen Brain Observatory.

A large fraction of functional interactions among NBG neurons involved significant negative CCG peaks among NBG neurons (nearly exclusively among pairs within areas, Table S2), and involved both putative excitatory and inhibitory neuron pairs (Tables S3), consistent with prior findings in mouse visual cortex (Senzai *et al*., 2019; Siegle *et al*., 2021) and other rodent neocortical areas (Fujisawa et al., 2008). Again, these enhanced functional interactions among NBG neurons were not simply explained by high firing rates: the probability of functional interactions among NBG neurons was markedly and significantly greater than among non-NBG neurons with high firing rates (those with rates greater than the mean of NBG neurons in the same area; Table S3).

Thus far we have examined NBG oscillations and spike timing during spontaneous activity – how does this relate to visual selectivity? Surprisingly, we found that NBG LGN neurons with ON versus OFF preferences showed clearly distinct NBG firing phase preferences, and these phase relationships were preserved across the visual cortical hierarchy. We first determined LGN neuron visual response preferences for luminance increments versus decrements (white versus black full screen flashes from grey background; Fig. 4A). As expected, many NBG LGN neurons showed clear onset responses to luminance increments (ON cells, n = 147; Fig. 4B), while others showed clear onset responses to luminance decrements (OFF cells, n = 77; Fig. 4C; see Methods for ON/OFF classification criteria). We then determined the preferred NBG firing phase (again during spontaneous activity) for ON and OFF neurons by constructing CCGs of every NBG neuron relative to the same reference cell within session (the ON cell with strongest NBG ACG power; see Methods). In this way, the strongest NBG ON cell defined zero phase within and across sessions. We plotted the relationship between visually evoked ON/OFF index for all neurons (no selection criteria) versus their preferred NBG firing phase during spontaneous activity and found a strong and significant correlation between the two measures (Pearson r = -0.55; *p* < *1e-31*; Fig. 4D). We then focused on the clearly ON versus OFF dominant neurons (same as in Fig. 3B, C) and investigated how these fired relative to NBG phase. ON versus OFF neurons showed concentrated and clearly separated NBG phase preferences (Fig. 4F), with a phase offset of π/2 (i.e., a difference of ∼ 5ms for NBG at 56 Hz). We then wondered if this preferred phase for NBG ON and OFF spiking in LGN was also maintained relative to NBG in V1 and HVAs. As before, we plotted NBG LGN neuron spike time histograms relative to the NBG LFP phase within cortical areas (cycle histograms, see Fig. 2E) and found that NBG ON and OFF neurons showed clear and distinct phase preferences relative to NBG LFP in V1 (Fig. 4G); remarkably, this phase separation of ON and OFF spiking was preserved relative to NBG LFP across the hierarchy of HVAs (Fig. S3). We then determined if an LGN neuron’s preferred NBG phase during spontaneous activity by itself could predict its ON/OFF visual selectivity (see Methods). Indeed, the preferred NBG phase of individual LGN neurons (relative to NBG within LGN) accurately identified their ON vs OFF visual selectivity (d’ = 2.2 ± 0.06; Fig. 4H); remarkably, visual ON/OFF selectivity of LGN neurons was also predictable solely from their preferred NBG firing phase relative to LFP in V1, LM, RL, AL, and AM (Fig. 4H), with predictions approaching chance level only in PM. Further, this significant predictability of LGN ON versus OFF channels from spontaneous NBG LFP showed a strong and significant correlation with the anatomical hierarchy (Fig. 4H; Pearson r = - 0.8; *p = 0*.*03*), just as was observed for the overall strength of NBG interactions from LGN across the visual cortical hierarchy (Fig. 2). Importantly, preferred NBG phase in LGN neurons relative to cortical LFP remained highly consistent throughout the duration of the recording (phase SD = ±0.65 radians, or ±1.72 ms for NBG at 60Hz; computed from 100 non-overlapping segments of 100s each in a single session across LGN and V1). This suggests that LGN ON versus OFF channels maintain a consistent, millisecond timescale phase offset relative to NBG activity in the input layers across V1 and HVAs.

**Figure 4.**
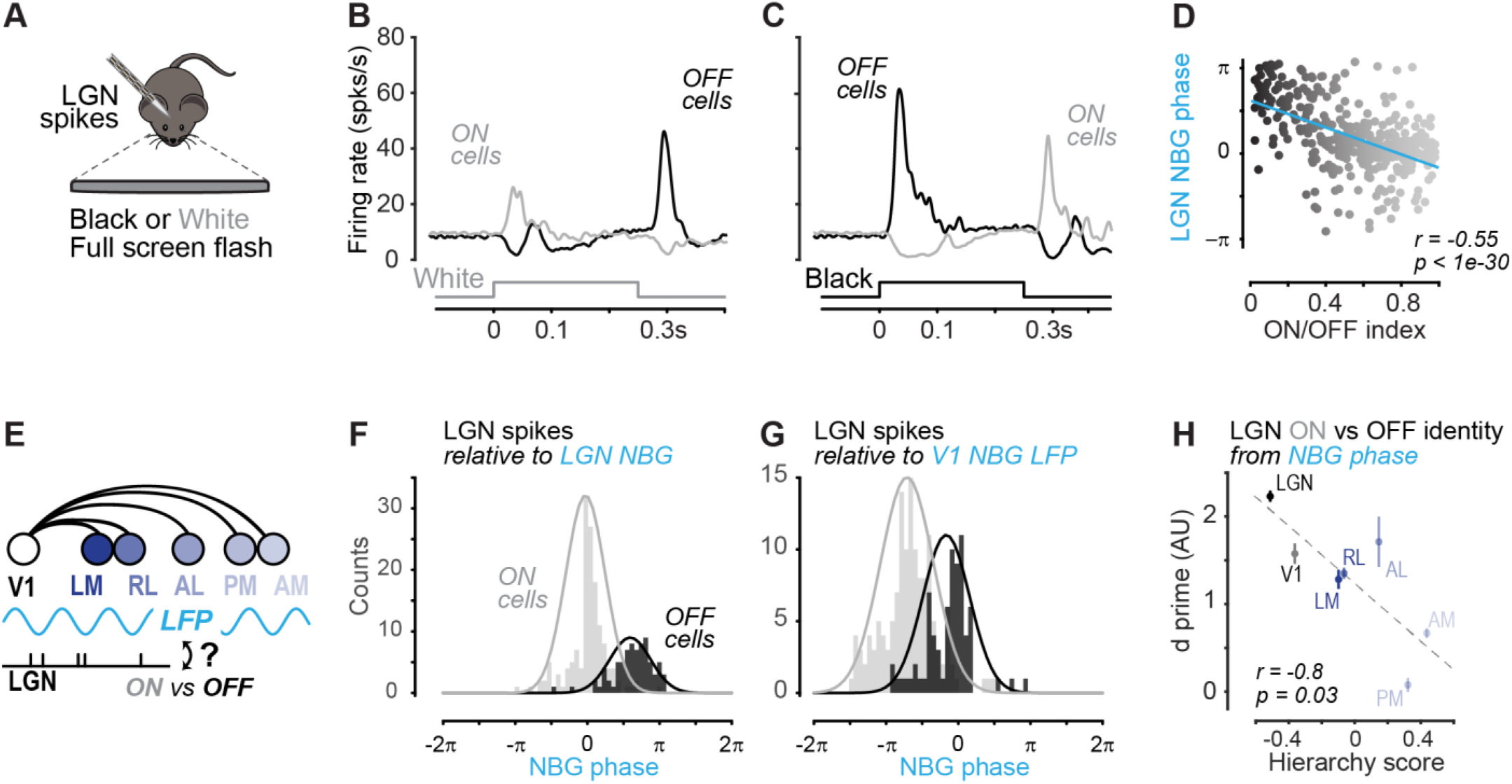
ON / OFF neurons in LGN spike at distinct NBG phases across LGN, V1, and HVAs. **A**. LGN spike responses measured during black or white full screen flashes. **B**. PSTH of LGN ON (grey, n = 147) and OFF (black, n = 77) responses to white stimulus across recordings (n = 14). See Methods for ON/OFF index calculation. **C**. Same neurons as B, in response to black stimulus. **D**. Scatter plot of ON/OFF index (abscissa) versus NBG neuron preferred phase (ordinate) for all ON/OFF LGN neurons (n = 397, 14 sessions). LGN NBG phase calculated relative to strongest NBG LGN ON neuron ACG (see Methods). Significant correlation between ON/OFF preference and NBG phase preference (Pearson rho = -0.55; *p* < *1e-31*). **E**. LGN ON and OFF neuron spikes were aligned to NBG LFP phase across V1 and HVAs. **F**. NBG phase of ON and OFF preference LGN neurons (same as B-D), relative to strongest NBG LGN ON neuron (see Methods). Gaussian fits to ON/OFF cell phase histograms show clear preferred phase clustering and separation. **G**. Same as F, but LGN spikes referenced to simultaneously recorded V1 L4 LFP. NBG phase is identified with spike-LFP cycle histogram (Fig. 2E). **H**. LGN ON versus OFF neuron preferences are predictable from LGN NBG spike phase relative to cortical LFP. Prediction accuracy (ordinate, d’) measured by calculating the distance between means of phase preference Gaussian fits (F and G) divided by the average SD of two fits (see Fig. S3). Discriminability (d’) of ON/OFF neurons by NBG phase shows significant correlation with hierarchy, and d’ remains above chance level for all areas except PM. All recordings in B – D, F – H from Allen Brain Observatory.

Finally, the major results shown here were unaffected when we controlled for potential confounds of volume conduction of LFPs, or for effects from mis-classifying NBG neurons. First, there was no dependence of HVA NBG power upon physical proximity to V1 (r = -0.04, *p = 0*.*8*; Fig. S4), while there was a clear and significant relationship between NBG power and HVAs according to their position in the visual hierarchy (r = -0.4, *p* < *2e-3*). Second, we recomputed all results using only LGN neurons with significant NBG ACGs: there remained a clear and significant relationship between visual cortical LFP and NBG LGN spiking, this varied as a function of anatomical hierarchy, and ON and OFF neurons in LGN were identifiable solely from their preferred NBG phase (Fig. S4).

## Discussion

Here we showed that narrowband gamma (NBG) activity synchronizes and spreads throughout the mouse thalamocortical visual system. NBG activity varied hierarchically from early to late stages of cortical processing. Spikes in LGN phase locked to NBG oscillations of LFP in V1, and throughout multiple higher visual areas (HVAs). NBG in multiple HVAs depended upon retinotopically aligned activity from V1. Neurons displaying NBG spiking formed distinct subnetworks with enhanced short latency functional connectivity within and across areas. Remarkably, LGN neurons that preferred luminance increments versus decrements (ON versus OFF pathways) showed distinct spike timing relative to NBG oscillations in LGN, V1, and HVAs. Taken together, these results identify synchronized NBG oscillations as a potential substrate for temporal coordination of spiking in functionally distinct subnetworks of neurons during awake vision.

We revealed that spontaneous NBG activity synchronized across LGN, V1, and HVAs. NBG activity was apparent at multiple scales from population level LFP oscillations to pairwise correlated spiking. Prior work in awake mice has investigated NBG activity in V1 (Niell and Stryker, 2010; Senzai *et al*., 2019; Speed *et al*., 2019), between LGN and V1 (McAfee et al., 2018; Saleem *et al*., 2017), and we now establish that NBG also invades multiple HVAs and synchronizes with NBG spiking in LGN. This spiking was phase locked to NBG LFP in the visual input layers of HVAs, echoing findings that NBG is strongest in L4 of V1 (Saleem *et al*., 2017; Speed *et al*., 2019). However, we also found significant NBG spiking and functional interactions among neurons in all layers of V1 and HVAs, consistent with spread of NBG beyond input layers (Speed *et al*., 2019). The areal and laminar distribution of NBG activity and its relationship to LGN and higher order thalamic projections forms an important topic for future study (Bennett et al., 2019; Blot et al., 2021; Harris *et al*., 2019); it will also be important to determine if NBG LGN neurons receive preferential input from distinct retinal ganglion cells (Sanes and Masland, 2015). Taken together, these findings identify that thalamocortical NBG oscillations could play the role of a “central clock” to reliably coordinate and synchronize spiking across the mouse visual system.

We found NBG oscillations engaged specific subnetworks of single neurons, and these displayed enhanced functional interactions among one another. These findings relied on examination of hundreds of thousands of simultaneously recorded neuron pairs in a unique public dataset of high-density, multi-site Neuropixels probes in multiple visual areas (Allen Brain Observatory, 2019; Siegle *et al*., 2021). In all structures examined, nearby neurons showed heterogeneity in the strength of NBG spiking. However, cross correlations among neurons with significant NBG activity showed greater likelihood for short-latency functional interactions (and common input) than neurons lacking NBG activity, even considering those with high firing rates (de la Rocha *et al*., 2007). It is important to note that short-latency peaks in cross correlations among LGN NBG neurons likely reflect common input, since recurrent excitatory connections are sparse in LGN, unlike in cortex. In V1 and HVAs, we found that many NBG pairs with significant functional interactions involved pairs of regular spiking (RS) and fast spiking (FS) neurons. These observations differ from prior findings showing strong NBG activity in excitatory but not inhibitory synaptic currents in excitatory neurons in V1 (Saleem *et al*., 2017). This could be explained by the enhanced detectability of functional interactions in cross correlations using spike trains from FS putative inhibitory interneurons that are highly active, as in prior reports (Fujisawa *et al*., 2008; Senzai *et al*., 2019). Even if excitatory neurons provide the dominant drive for NBG, high-density sampling with Neuropixels probes could facilitate isolation of spikes and interactions between FS and RS neurons compared to prior studies that used less dense electrodes. It will be important to determine how NBG activity engages the diversity of cortical cell types in V1 and HVAs (Veit et al., 2017), and how subthreshold excitation or inhibition influences NBG spiking activity.

We found that the strength of NBG activity depended upon retinotopy and obeyed the anatomical hierarchy of visual areas. The simplest interpretation for hierarchical NBG activity is that it merely reflects the density of feedforward visual connectivity from LGN to V1 to HVAs. Our findings that V1 inactivation specifically reduced NBG power in multiple downstream HVAs in a retinotopic manner is consistent with this scheme, and suggests that neurons across visual areas with shared spatial receptive fields also share NBG activity. Since NBG in L4 of V1 promotes subsequent detection of spatially localized visual stimuli (Speed *et al*., 2019), a testable prediction from our findings is that increased pre-stimulus NBG power provides an effective state to synchronize stimulus-evoked spikes in retinotopically aligned neurons across LGN, V1, and HVAs, leading to improved visual perception.

How might synchronized NBG oscillations underlie improved perception of visual stimuli? Although our results focused on spontaneous activity, a potential clue is offered from the unexpected link between spontaneous NBG firing phase in ON and OFF pathways. In LGN, ON and OFF neurons fired at distinct phases of NBG, and this phase separation was maintained relative to NBG LFP in V1 and HVAs, particularly in areas RL, LM, and AL. These specific HVAs are necessary for detection of stimulus contrast (Goldbach et al., 2021; Jin and Glickfeld, 2020). Synchronized pre-stimulus NBG activity could dictate the timing and propagation of the very first spikes in cortex that are essential for perception (Resulaj et al., 2018). Indeed, in retina, first spike latencies in ON and OFF pathways carry the most information about the visual scene (Gollisch and Meister, 2008), and this may precisely shape spike timing in LGN (Koepsell et al., 2009; Storchi *et al*., 2017). Even if strong visual stimulation transiently suppresses NBG (Saleem *et al*., 2017), precisely timed activation then silence may efficiently encode visual information across neural populations (Schneidman et al., 2011). Thus, synchronization and propagation of a narrowband, phase-locked, stimulus-independent oscillation by LGN throughout visual cortical areas could provide a substrate for temporal coding in ON and OFF pathways throughout the mouse visual system. This establishes a novel testbed for resolving long-standing and open questions regarding oscillations, temporal coding, and alternative modes of communication underlying visual perception (Fries, 2015; Gray, 1999; Kohn *et al*., 2020; Shadlen and Movshon, 1999).

## Acknowledgements

We thank members of the Haider lab and Aman Saleem for feedback. This work was supported by the Alfred P. Sloan Foundation’s Minority Ph.D. (MPHD) Program Fellowship (to J.D.R.), the Whitehall Foundation (to B.H.), the Alfred P. Sloan Foundation Fellowship In Neuroscience (to B.H.), National Institutes of Health Neurological Disorders and Stroke (NS107968 to B.H.), National Institutes of Health BRAIN Initiative (NS109978 to B.H.), and by a grant from the Simons Foundation (SFARI 600343, B.H.)

## Author Contributions

D.S. accessed the database, wrote analysis code, and analysed all experimental data; J.D.R., K.P. performed silicon probe plus optogenetics experiments; D.S., B.H. wrote the manuscript with feedback from all authors.

## Inclusion and Diversity Statement

We worked to ensure sex balance in the selection of non-human subjects. One or more of the authors of this paper self-identifies as an underrepresented ethnic minority in science. One or more of the authors of this paper received support from a program designed to increase minority representation in science. The author list of this paper includes contributors from the location where the research was conducted who participated in the data collection, design, analysis, and/or interpretation of the work.

## Methods

### Resource Availability

#### Lead Contact

All requests for resources should be directed to and will be fulfilled by Bilal Haider.

#### Materials Availability

This study did not generate new unique reagents or materials.

#### Data and Code Availability

All data structures and code that generated each figure will be deposited on Figshare (https://doi.org/10.6084/m9.figshare.19666314) and linked from the corresponding author’s institutional webpage upon publication.

### Experimental Model and Subject Details

All procedures were approved by the Allen Institute’s Institutional Animal Care and Use Committee and Institutional Animal Care and Use Committee at the Georgia Institute of Technology.

#### Experimental subjects - Allen Brain Observatory

Recordings from the Allen Brain Observatory – Visual Coding database are fully detailed elsewhere (Allen Brain Observatory, 2019; Siegle *et al*., 2021). 45 male and 13 female mice were used. 32 of 58 subjects had single unit recordings in LGN; we focused on 14 / 32 that had ≥10 NBG LGN neurons recorded simultaneously. In the 14 sessions with ≥10 NBG LGN neurons, we analyzed 316 ± 59 cells recorded simultaneously (all areas) and 51,404 ± 19,331 possible pairwise interactions per experiment. The Dataset for this study was last accessed and compiled on December 30, 2020.

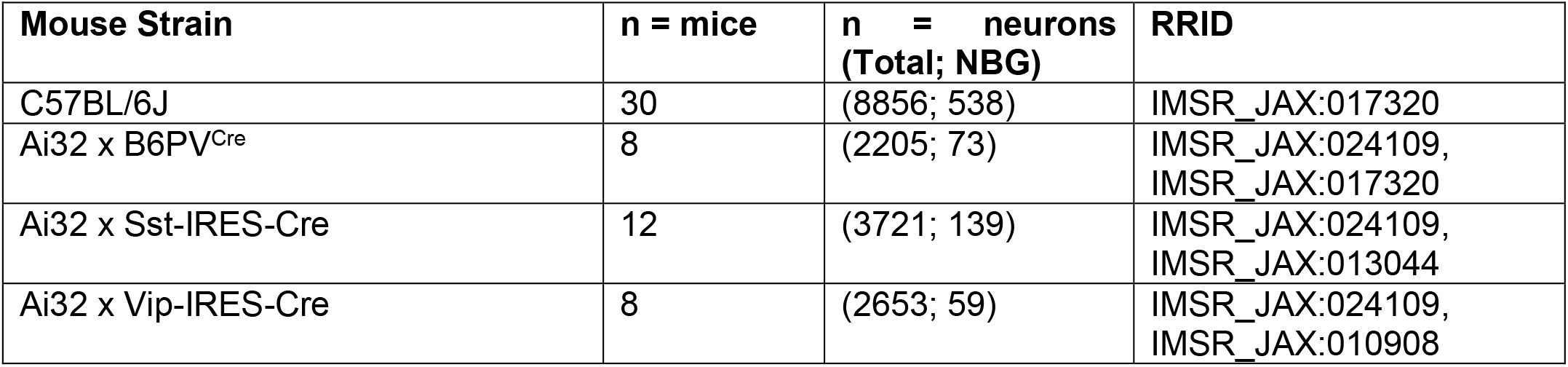

#### Experimental subjects - Haider lab

Detailed methods for neural recording and optogenetic inactivation have been described previously (Speed *et al*., 2019; Speed et al., 2020). Mice (5 – 8 weeks old; reverse light cycle individual housing; bred in house) were chronically implanted with a stainless steel headplate with a recording chamber during isoflurane (1-2%) anesthesia. After implant surgery mice recovered for 3 days before experimentation. Recordings in Haider lab used male and female mice.

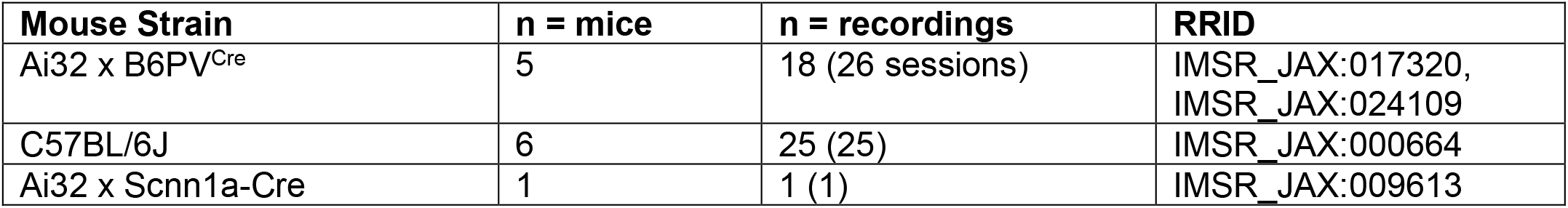

| Mouse Strain | n = mice | n = recordings | RRID |
| --- | --- | --- | --- |
| Ai32 x B6PV<sup>Cre</sup> | 5 | 18 (26 sessions) | IMSR_JAX:017320,<br>IMSR_JAX:024109 |
| C57BL/6J | 6 | 25 (25) | IMSR_JAX:000664 |
| Ai32 x Scnn1a-Cre | 1 | 1 (1) | IMSR_JAX:009613 |

Across all datasets, there was no allocation strategy for selecting subjects, since there were no comparisons between experimental groups of separate subjects.

## Method Details

### Recordings

#### LGN, V1, HVA Neuropixels recordings – Allen Brain Observatory

Detailed recording procedures are described elsewhere (Allen Brain Observatory, 2019; Siegle *et al*., 2021). Data was retrieved from the database using their proprietary software development kit (SDK). Data was compiled last on December 20, 2020. Detailed instructions for accessing the database and recording sessions with analysis code will be publicly deposited and linked from the lead contact’s institutional website.

#### V1 inactivation and HVA recordings – Haider lab

All procedures were approved by the Georgia Institute of Technology Institutional Animal Care and Use Committee (IACUC). A custom-built stainless steel head post with a recording chamber (11 mm inner diameter) was lightly affixed to the skull using veterinary adhesive (VetBond). Following headplate fixation, a glass coverslip (5 mm diameter, #1 thickness ∼0.15 mm) was centred over the representation of V1 and HVAs (centre of window at ∼ 2.4 mm lateral to midline and ∼ 2.4 mm anterior to lambda) and bonded to the skull using VetBond. Mice were individually housed and monitored for full recovery for at least 3 days before intrinsic signal imaging (ISI) of V1 and HVAs, as detailed in our prior studies (Nsiangani et al., 2022). Briefly, ISI was performed during isoflurane anesthesia with sedation. The cortex was illuminated with green or red light to capture hemodynamic or blood oxygenation signals, respectively. Custom visual stimulation and image acquisition systems computed retinotopic maps of stimulus selectivity for azimuth and elevation, and these were used to compute visual field sign maps that defined boundaries and extent of V1 and the HVAs, consistent with established methods (Juavinett et al., 2017; Zhuang et al., 2017). Following ISI, mice were habituated to head fixation for 3 to 5 days before undergoing awake recordings as in prior studies (Speed *et al*., 2019). On recording days, small craniotomies (∼100-500 μm) were made in HVAs under isoflurane anesthesia, using the ISI maps and vasculature as references (Nsiangani *et al*., 2022). The skull overlying two retinotopically distant sites in V1 (∼ 60 to 90° apart in azimuth) was thinned to facilitate optical stimulation. Mice were allowed to recover for >3 h, and then acute awake recordings were made with multi-site silicon probes (Neuronexus; A 1×32) spanning all layers of the cortex. Electrodes were advanced ∼1000 μm below the cortical surface. The signals were acquired at 30 kHz (Blackrock Microsystems) and filtered at 0.3 – 300 Hz to acquire the LFP signal. LFP power was analyzed on the channel with greatest raw NBG power, typically in L4. Experiments reported here only examined long periods of spontaneous activity (no visual stimulation). V1 was locally silenced by activating ChR2 expressed in parvalbumin inhibitory neurons as in our prior studies (Speed *et al*., 2019; Speed *et al*., 2020). Laser stimulation (6.5mW power, 1s duration with 0.1s onset and offset ramps, inter-trial interval duration randomly selected from 1 – 6s per trial) was confined to a circular spot (∼0.2 mm diameter half-width, measured with a beam profiler) using custom optics, and the spot was targeted to portions of V1 that matched / mismatched the retinotopic coordinates of the craniotomies targeted to HVAs (using ISI maps for azimuth and elevation; Fig. S3). Laser stimulation was only delivered on 25-33% of randomly selected trials with the remaining trials serving as the interleaved control trials. At the conclusion of optogenetic experiments, retinotopy was confirmed at the HVA recording sites by presenting briefly flashed bright and dark bars throughout the visual field (Nsiangani *et al*., 2022). Retinotopy of the thinned V1 sites was also subsequently electrophysiologically confirmed the same way. Using azimuth coordinates from both ISI maps and the preferred azimuth of the LFP responses recorded in HVAs, we classified V1 and HVAs retinotopy as “matched” when they were both located within the binocular visual field (0 to 40° azimuth), or both located in the monocular visual field (>55° azimuth). For matched conditions, the V1 and HVA RFs were 9.6 ± 2.6° apart (mean ± SEM), within the average width of single neuron spiking RFs in mouse V1 (Niell and Stryker, 2008). “Mismatched” experiments paired recordings and stimulation across binocular and monocular locations: here, V1 and HVA RFs were 46.8 ± 3.2° apart, significantly more separated than the matched experiments (p = *2*.*2e-04*, Wilcoxon rank sum test). Importantly, matched versus mismatched experiments varied only in azimuthal (horizontal) coordinates: differences in elevation (vertical) retinotopy between V1 and HVA sites were small and comparable across groups (Matched: 10.0 ± 1.3° apart; Mismatched 8.3 ± 1.3° apart; *p = 0*.*32*, Wilcoxon rank sum test; coordinates assigned from ISI maps).

#### LGN recordings – Haider lab

Detailed methods have been described previously (Williams et al., 2021). Briefly, LGN was targeted with stereotaxic coordinates (−2.5 mm posterior to bregma -2.5 mm lateral from midline). Mice were habituated to the recording environment prior to awake recordings. Single shank electrodes (Neuropixels 1.0 IMEC) were used to record from LGN (n = 23 experiments). Each probe contains 960 channels, of which a subset of 383 were used for recording. Spikes were acquired at 30 kHz and LFP at 2.5 kHz via a PXIe card, National Instruments board, and Spike GLX software. In a subset of experiments (n=3), single shank electrodes (NeuroNexus, A1×32 Poly 3) were used to record from LGN and signals were acquired through a Cereplex Direct (Blackrock Microsystems). In all experiments, visual stimuli were shown and unit responses monitored on-line as electrodes advanced to LGN (∼ 3.0-3.2 mm below the dura), and in most instances confirmed with histology. Spatial receptive fields were mapped with black and white squares to confirm functional responses consistent with LGN neurons.

### Visual stimuli

#### Visual stimuli – Allen Brain Observatory

Detailed stimulus parameters are described elsewhere (Allen Brain Observatory, 2019). We analyzed responses to full-field flashes of white and black (0.25 s duration, 2 s inter-trial interval with uniform grey screen) to categorize ON and OFF preference neurons in LGN.

### Quantification and Statistical Analysis

#### Spike sorting

##### Spike sorting – Allen Brain Observatory

Neurons were pre-sorted and packaged with several pre-computed quality metrics, as detailed elsewhere (Allen Brain Observatory, 2019). We plotted histograms of spike waveform widths and observed a clear bimodal distribution in cortical recordings with a partition at 0.42 ms that separated regular spiking (RS) and fast spiking (FS) groups. LGN recordings showed unimodal spike waveform width histograms, with a large majority having spike widths > 0.38 ms (91%; 1,187 / 1,306), including the majority of NBG neurons (91%; 415 / 455). A minority of LGN neurons showed spike widths between 0.2 – 0.38 ms (9%, 119 / 1,306), including just 8% of NBG neurons (40 / 455). We interpret this to suggest that the great majority of NBG LGN neurons correspond to classically described excitatory thalamocortical relay neurons (Guido, 2018; Williams et al., 1996).

#### Visual response analysis

##### Laminar identification – Allen Brain Observatory

The earliest sink of stimulus triggered current source density (CSD) in V1 corresponds to the site of strongest LGN input in layer 4 (Lien and Scanziani, 2013; Speed et al., 2019; Speed et al., 2020). This anatomical relationship is less clearly defined for CSDs in HVAs (RL, LM, AL, AM, PM). For consistency, we analyzed NBG power at the earliest stimulus triggered CSD sink channel across V1 and HVAs. All cortical channels were restricted to the hundred channels below the topmost channel expressing electrical activity (1mm total). We defined the earliest sink channel as the one that reached the half-maximum amplitude of the earliest sink with shortest latency (Fig. S2). This analysis identifies the site of the earliest functional visual input across all areas but cannot clearly identify the anatomical source of this input in HVAs.

#### NBG LFP power analysis

We quantified residual NBG power of LFP by modifying previously established methods (Saleem *et al*., 2017). We first fit a linear regression model on the logarithmic power (dB) of the frequency domain LFP (using fitlm.m function in MATLAB). To compute the residual NBG power, we first fit all data points in the frequency range spanning 30 – 90 Hz (excluding 50 – 70 Hz). We then defined the residual NBG power as the average LFP power between 50 – 70 Hz that remained after subtracting the power in the NBG range predicted from the smoothed linear fit. This method was used to calculate raw LFP residual NBG power and stLFP residual power. Residual NBG power was considered in excess when mean power in NBG frequency range was > 1 SD above the linear fit (using movmean.m and movstd.m from MATLAB with 1000 datapoints). As a control measure during optogenetic experiments (Fig. 2), we calculated residual broadband gamma power in the same manner by computing linear fits from 20 – 90 Hz (but excluding 30 – 70 Hz), then measuring residual broadband gamma power from 30 – 70 Hz, consistent with the range for broadband gamma in mouse V1 (Veit *et al*., 2017).

#### NBG neuron identification in spike cross-correlograms and autocorrelograms

##### NBG neuron identification with auto-correlogram

Spike-trains from cells in all visual areas were binned at 0.5ms resolution (‘histcount’ in MATLAB). Autocorrelograms (ACGs) and cross-correlograms (CCGs) were calculated with ‘xcorr’ in MATLAB.

The first step in NBG neuron identification examined ACGs. We computed frequency domain power of the “sidebands” of the ACG function (lags between 20ms to 120ms; 201 points, 0.5ms bin size) to avoid distortion caused by the central ±10 ms of the ACG (Fig. S1). NBG neurons identified with the ACG fulfilled either of the following conditions: *1)* Maximum power _[50–70Hz]_ > 0.9 * maximum power _[40-300Hz]_; *2)* Maximum power _[50–70Hz]_ is local maximum & Maximum power _[50–70Hz]_ > 0.9 * maximum power _[50-300Hz]_. These criteria were implemented to avoid exclusion of NBG neurons that also exhibited strong “classic” broadband gamma power near 40 Hz. We excluded any NBG neurons identified with the ACG if they did not form significant NBG CCGs with any other simultaneously recorded neuron (58 / 272; 21% of neurons). These exclusion criteria were implemented to remove any potentially non-physiological contamination of spike trains visible in the ACG, and likely lead to an underestimate of the true number of NBG neurons in the recordings.

##### NBG neuron identification with cross-correlogram (NBG-CCG)

The main findings of this study are based on NBG neuron identification using CCGs. Our reasoning for using CCGs is that low firing rate neurons or those with long refractory periods may not manifest ACGs with oscillatory firing across multiple successive NBG cycles (Fig. S1). It is important to note that CCGs likely reflect a mixture of direct functional interactions and indirect interactions due to common input; this is particularly true among cell types that are known to have little local connectivity, such as among LGN neurons. The CCG analysis conditions spike histograms on joint firing at NBG frequencies across neuron pairs and thus isolates a larger population of NBG neurons than solely ACG analysis (n = 400 identified with CCG analysis; n = 214 with ACG analysis; 14 sessions). Importantly, the identification of NBG neurons in CCGs was highly sensitive to spike timing controls (Fig. S1), and the main results are nearly unchanged when analyzing NBG neurons identified solely with ACGs (Fig. S4).

CCGs were calculated with the same resolution (0.5ms) as ACGs, but the central ±50ms of the CCG were analyzed in the frequency domain. CCGs were normalized to have maximum 1 and Fourier transformed. Both neurons of the pair that fulfilled all four of the following conditions were classified as NBG neurons: *1)* Magnitude _[60Hz]_ > Mean magnitude _[30–150Hz]_ + 2*SD _[30–150Hz]_; *2)* Magnitude _[60Hz]_ > Mean magnitude _[150–1kHz]_ + 2*SD _[150–1kHz]_; *3)* Magnitude _[60Hz]_ > 2* Max magnitude _[150–1kHz]_ ; *4)* NBG magnitude should be higher than 2 (unitless quantity, from normalized FFT).

We ensured that neurons that passed both ACG and CCG criteria were not double counted as NBG neurons, and if a neuron only passed the ACG criteria but formed no significant NBG CCGs with any simultaneously recorded neurons, it was excluded from further analysis.

##### Comparisons and controls for NBG neuron identification with CCG and ACG

We tested if the identification of NBG neurons using CCG power was erroneously caused by CCG power driven by neurons with strong NBG ACGs. We computed NBG CCG power using pairings of all simultaneously recorded neurons, with one group comprised of NBG neurons identified by CCG, and another comprised of NBG neurons identified by ACG power (Figure S1K). We assigned NBG CCG power of a neuron as the highest NBG power among CCGs computed from all simultaneously recorded neurons. The distributions of NBG power in neurons identified by CCG or ACG were largely overlapping, but significantly different due to the large number of samples (Fig. S1K). Next, even when we excluded CCGs containing neurons with significant NBG ACGs, the distributions of NBG power in CCGs were still largely overlapping (Fig. S1L). Classifying NBG neurons through CCGs — when excluding neurons with significant NBG ACGs — correctly classified 56 out of 186 NBG neurons identified through CCG criteria using all pairwise CCGs and classified 161 out of 214 NBG neurons identified through ACG criteria.

We estimated the probability of false identification of NBG neurons with CCGs by computing shuffled CCGs between NBG neurons identified across different recording sessions. We performed CCG between NBG-ACG cells in LGN in one session (n = 34 significant NBG ACG neurons) and all cells in a different session (n= 242 NBG neurons). Among these 8228 CCGs, none was falsely identified as a CCG indicating NBG neurons according to our criteria.

We also verified that identification of NBG neurons with CCGs within session depended upon NBG spike timing. We jittered spike times in a NBG neuron identified through CCG, and calculated the CCG between this jittered spike train and a spike train of a neuron with significant NBG ACG power. In this example pair, 0/10k jittered CCGs were identified as containing NBG neurons (Fig. S1F), and across 30 other pairs drawn randomly across experiments, only 1/300k jittered CCGs was erroneously identified as a NBG pair. These results show that false positive identification of NBG neurons in CCGs through “power leakage” of neurons with strong NBG spike ACGs is unlikely.

We also examined the features of NBG neurons in LGN identified with ACG criteria (versus CCG criteria) and found that these were overlapping but significantly different in several regards compared to NBG neurons classified with CCG criteria (Fig S1), including higher average firing rates (12.2 ± 4.1 vs 10.1 ± 2.9, median ± IQR) and greater amount of NBG power in CCGs (17.7 ± 2.1 vs 16.2 ± 2.6), even when excluding CCGs between pairs of neurons that both passed the ACG criteria (Fig. S1L). Nonetheless, as seen in Fig. S4, the main findings of the study were nearly identical even when only analyzing NBG neurons identified solely with single neuron spike ACG criteria.

##### Detection of functional connectivity in cross-correlograms

We detected functional connectivity between cells with previously established methods (Fujisawa *et al*., 2008; Senzai *et al*., 2019; Stark and Abeles, 2009) with a few modifications. First, we performed cross-correlation on pairs of spike-trains, then applied a bandstop filter between 50 – 70 Hz (ideal filter applied in frequency domain representation of the correlogram through fft.m function in MATLAB) to minimize spurious threshold crossings caused by strong NBG correlated firing. Second, we detected values in the filtered correlogram that deviated from the expected Poisson process significance threshold in a short time scale (1ms – 4.5ms for functional connectivity pairs, 0ms – 1ms for common input pairs). To compute the Poisson significance threshold, we convolved the filtered correlogram with the Gaussian filter (gausswin.m function in MATLAB) with SD = 7ms (total filter length 42ms at ± 3SD). The convolved correlogram approximates the hypothetical Poisson process where the spike count mean equals the variance and has been shown to detect significant deviations from the Poisson expectation at ∼ p ≤ 1 e10^−6^ threshold, as described previously (Senzai *et al*., 2019; Stark and Abeles, 2009). If more than two consecutive points fell above or below the Poisson significance threshold between 1.5ms to 4.5ms (including edges), the pair was identified as one exhibiting functional connectivity. We did not factor in transmission delays (e.g., from LGN to V1) into our acceptance window in order to maintain consistent and conservative criteria for functional interactions both within and across multiple visual areas (only 1 additional LGN-V1 pair would have been included with a larger acceptance window factoring in the delay). Correlograms fulfilling the above criteria in the positive direction were classified as excitatory interactions, and in the negative direction classified as inhibitory interactions (Table S2). Correlograms fulfilling the above criteria with peaks at 0ms – 1ms were classified as common input pairs. We only considered significant common input pairs with positive crossings of the significance threshold and excluded CCGs with non-physiological profiles (i.e., exceedingly high peaks in the central bin).

##### Spike-LFP cycle histogram analysis

We used two complementary methods to assess spike-LFP coherence: LFP cycle histograms (Colgin et al., 2009) and spike-triggered LFP. Cycle histograms were constructed by first bandpass filtering the LFP in the NBG range (50 – 70 Hz, bandpass.m function in MATLAB) and marking the location of the local maxima (findpeaks.m function in MATLAB). Then, we aligned the relative location of LGN spikes between the local maxima of the bandpass filtered LFP (spike phase; Equation below) Lastly, we normalized the histogram of these spike phases to have maximum amplitude 1 (using histogram.m function in MATLAB; binsize = 0.05 radian, from 0 to 2π)

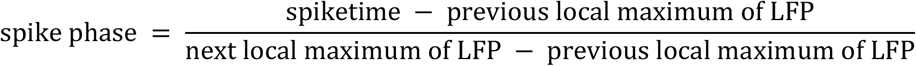

We computed three metrics from cycle histograms by fitting a cosine (by extracting the first Fourier component of the histogram using fft.m function in MATLAB). The Cycle SNR (Fig. 2F) is the power ratio of fitted cosine and the residual (using snr.m function in MATLAB). Cycle phase is the phase of the fitted cosine (using angle.m function in MATLAB).

##### Spike triggered LFP analysis

Spike triggered LFP (stLFP) was computed as follows:

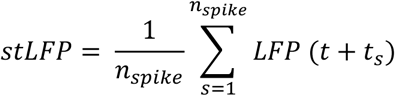

*n*_*spike*_ is the total number of spikes of a neuron used for triggering LFP vectors. stLFP vectors from raw LFP at the earliest current sink channel (identified from CSD analysis; Fig. S2) were extracted surrounding the spike time from [-0.2s, 0.25s]. The time step *t* was 8ms (i.e., LFP vector sampled at 1250 Hz), consistent with prior work on stLFPs (Telenczuk et al., 2017).

##### Correlation of NBG activity and visual hierarchy analysis

We computed correlation coefficients between the above-mentioned metrics and the anatomical hierarchy scores. Correlation coefficients (Pearson) and p-values were calculated with the corr.m function in MATLAB. Anatomical hierarchy scores of visual areas were taken from prior Allen Brain studies (Harris *et al*., 2019; Siegle *et al*., 2021). Numerical scores were extracted from Figure 3. C, F, I, L in (Siegle *et al*., 2021) using GRABIT.m (MATLAB Central File Exchange).

##### Correlation of NBG activity in HVAs versus distance to V1 (control for volume conduction)

To control for the potential influences of LFP volume conduction (Kajikawa and Schroeder, 2011) from V1 to HVAs, we assessed the relationship between stLFP power (and cycle histogram SNR) in HVAs as a function of distance from V1. Here, we analyzed a subset of recordings in the Allen Brain Observatory dataset: those with ≥10 LGN NBG neurons, clear LFP / CSD in V1, and all V1 and HVA recording locations verified with the Allen Mouse Brain Common Coordinate Framework (CCF; see https://atlas.brain-map.org/). In this subset (12 out of 58 total sessions), 12/12 recordings had probes in V1, 10/12 in RL, 6/12 in LM, 8/12 in AL, 7/12 in PM, and 10/12 in AM. We computed the distance between the earliest sink channel in each HVA (see Fig. S2) relative to the earliest sink channel of V1 (Layer 4). We then assessed correlations between the stLFP and cycle histogram SNR for HVAs (as in Fig. 2) relative to distance of HVAs from the V1 NBG source. The stLFP and cycle histogram SNR in HVAs was poorly correlated with distance to V1 (Fig. S4). Instead, stLFP and cycle histogram SNR were both strongly and significantly correlated with the functional hierarchy position of HVAs (Fig. 2).

##### Conditional probability analysis

We analyzed the conditional probability of functional connectivity between NBG pairs, and pairs formed by non-NBG neurons (Fig. 3). The analysis consists of counting the number of functional connectivity pairs among cells of interest then dividing by the total number of possible pairs among cells of interest. For the total number of possible pairs involving short-latency lagged interactions (1 – 4.5ms), the total possible connectivity among pairs was bidirectional n*(n-1); for the total possible number of common input pairs (0 to 1ms lagged), within area connectivity is not bidirectional among elements so was calculated as n*(n-1) / 2. Common input pairs across areas were bidirectional and the total possible connectivity among pairs was calculated as n*(n-1).

Then, we computed significance using a binomial t-test with the formula below:

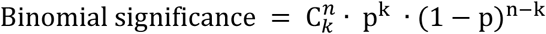

where n is total number of possible pairs, C denotes combination, k is number of functional connectivity (or common input) pairs, and p denotes the null hypothesis probability. This null hypothesis was computed as the probability of functional connectivity among non-NBG neurons (Fig. 3), or among non-NBG neurons with high firing rates (Results). Since correlations increase with firing rates (de la Rocha *et al*., 2007) this provides a more stringent “upper bound” null hypothesis for assessing NBG neuron functional interactions. In each area under consideration, we classified non-NBG neurons as “high firing” if their rates were greater than the mean rates of NBG neurons exhibiting significant CCGs in that same area. These null hypothesis estimates found P(functional connectivity | non-NBG & high firing rate) as 0.02% (LGN to V1), 2.7% (V1 to V1), and 0.01% (LGN to HVA).

We also tested the probability of functional connectivity among the “network” of ON or OFF dominant NBG neurons from LGN to HVAs. We found that there was no significant difference for LGN ON or OFF neurons to show preferred functional interactions with ON or OFF neurons in V1 and HVAs (defined by their responses to full screen flash stimuli, described below). That is, NBG ON and OFF neurons in LGN showed equal probabilities of functional interactions with both ON and OFF neurons in downstream cortical areas (*p = 0*.*7*, Wilcoxon rank sum tests), as expected since most cortical neurons typically show a broad range of ON/OFF selectivity (Williams *et al*., 2021).

##### ON/OFF classification and analysis in LGN

NBG neurons in LGN were classified as ON or OFF preference cells from the transient response to luminance increments and decrements during a full screen flash stimulus. Each cell had a total of 150 trials of stimulus responses to both luminance increments (ON; grey to white and black to grey) and decrement. (OFF; grey to black and white to grey). The number of spikes transiently responsive to the luminance change (25 – 75ms after the stimulus) were counted and used to compute ON/OFF index.

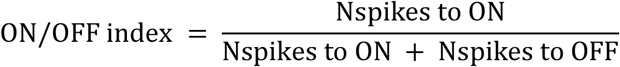

Cells with ON/OFF index > 0.7 were identified as ON dominant cells and those < 0.3 as OFF dominant cells, similar to prior studies (Schroder et al., 2020). Poorly responsive NBG LGN cells (evoked response < 2.7 spikes/sec) were excluded. Out of 400 neurons, 147 cells were identified as ON dominant and 77 as OFF dominant (14 recording sessions).

##### NBG phase analysis

We computed NBG phase of each NBG neuron in LGN in two different ways: through cortical LFP triggered cycle-histograms (described above in *‘Spike-LFP cycle histogram analysis’*), and through spike cross-correlation analysis. For this, we performed cross-correlations (0.5ms bin; window ±50ms) among all simultaneously recorded NBG LGN neurons. We computed CCGs between each NBG LGN neuron and one reference NBG neuron in the session (the ON cell with strongest NBG power in its ACG within the session). We bandpass filtered the correlogram in the NBG range (50 -70Hz; bandpass.m function in MATLAB). NBG phase was defined as the Hilbert phase at 0ms lag. (hilbert.m and angle.m functions in MATLAB). We estimated the mean and standard deviation of ON/OFF NBG phase histograms within each area by fitting Gaussian functions (Fig S3). We first normalized each ON/OFF NBG phase histogram by transforming the range of phase from [-π to π] to [(circular mean -π), (circular mean+π)] to account for circular phase. Circular mean was calculated using meanangle.m (MATLAB Central File exchange). Then we fit Gaussian distributions to the histograms using fitdist.m function in MATLAB with “normal” option.

---

**Figure S1.**
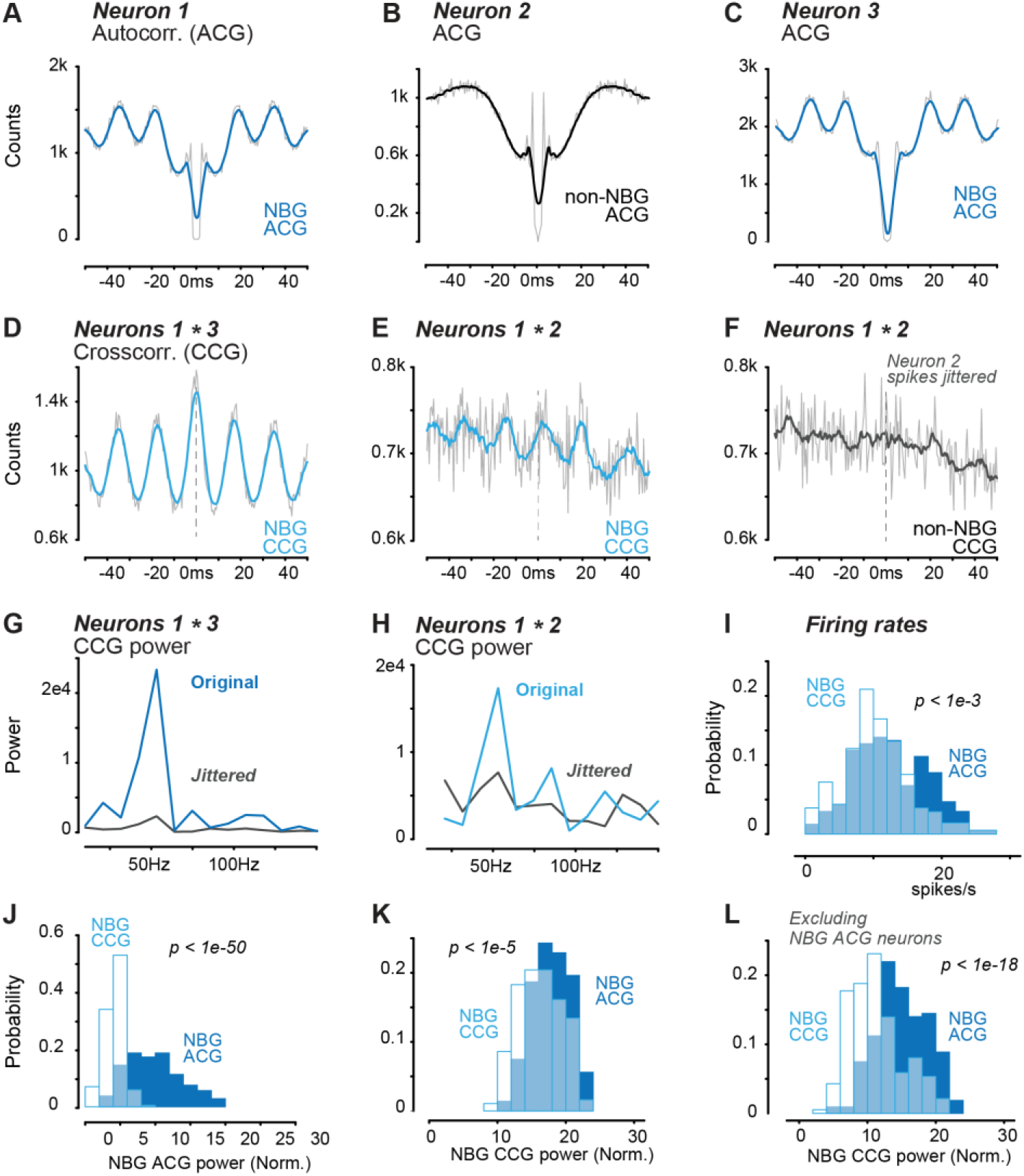
Auto-correlogram (ACG) versus cross-correlogram identification of NBG neurons in LGN. **A**. Auto-correlogram of example NBG neuron in LGN (112k spikes in 170 min of recording, 0.5ms bins; grey: raw ACG, blue: ACG of 10 point moving average, same throughout A-F). **B**. Same recording as A, ACG of example NBG neuron in LGN identified by CCG but not ACG criteria (111k spikes). **C**. Same recording as A, ACG of another NBG neuron in LGN (162k spikes). **D**. CCG of NBG neurons 1 and 3 (in A and C). The CCG power also classifies both as NBG neurons (see Methods for criteria). **E**. CCG of NBG neuron 1 and non-NBG neuron 2 (classified by ACG power). The CCG power classifies both as NBG neurons. **F**. Same neuron pairs as in E, but with jittered spike train of neuron 2 (uniformly distributed noise drawn from [-20ms, 20ms] interval added to every spike time). Out of 10k CCGs with jittered spike trains in neuron 2, none show classification of neuron 2 as a NBG neuron, indicating precise spike timing in neuron 2 relative to neuron 1 drives NBG CCG. **G**. Power spectrum of original (blue, same as D) and jittered (light blue) CCGs between neurons 1 and 3. NBG power decreased by 9.9 dB after jittering spikes. **H**. Power spectrum of original (blue, same as E) and jittered (light blue, same as F) CCGs between neurons 1 and 2. NBG power decreased by 3.6 dB after jittering spikes. **I**. Histogram comparing firing rates in NBG neurons defined by CCG (open bars; 10.1 ± 2.9 spikes/s, median ± IQR/2, n = 186 neurons in 14 sessions) versus ACG (filled bars; 12.2 ± 4.1 spikes/s; n = 214 neurons in 14 sessions *p = 1*.*03e*^*-4*^, Wilcoxon rank sum test). Same neurons shown in I – L. **J**. Histogram comparing NBG power in ACG of neurons identified with CCG power only (open bars; 4.3 ± 0.8; median ± IQR/2) and neurons identified solely with ACG power (filled; 9.3 ± 2.9; *p = 4*.*0e*^*-53*^, Wilcoxon rank sum test). **K**. Histogram comparing normalized NBG power of neurons classified from CCG (open bars; 16.2 ± 2.6; median ± IQR/2) versus ACG (filled; 17.7 ± 2.1; *p = 4*.*8e*^*-6*^, Wilcoxon rank sum test). For each neuron, the maximum NBG CCG power among all other simultaneously recorded neurons was plotted. NBG Power normalized to broadband power (40 -300Hz). **L**. As in K, but excluding CCGs with NBG ACG neurons from pairwise CCGs. NBG CCG power 10.5 ± 2.5 (median ± IQR); NBG ACG power 14.4 ± 2.8; *p = 1*.*1e*^*-19*^, Wilcoxon rank sum test).

**Figure S2.**
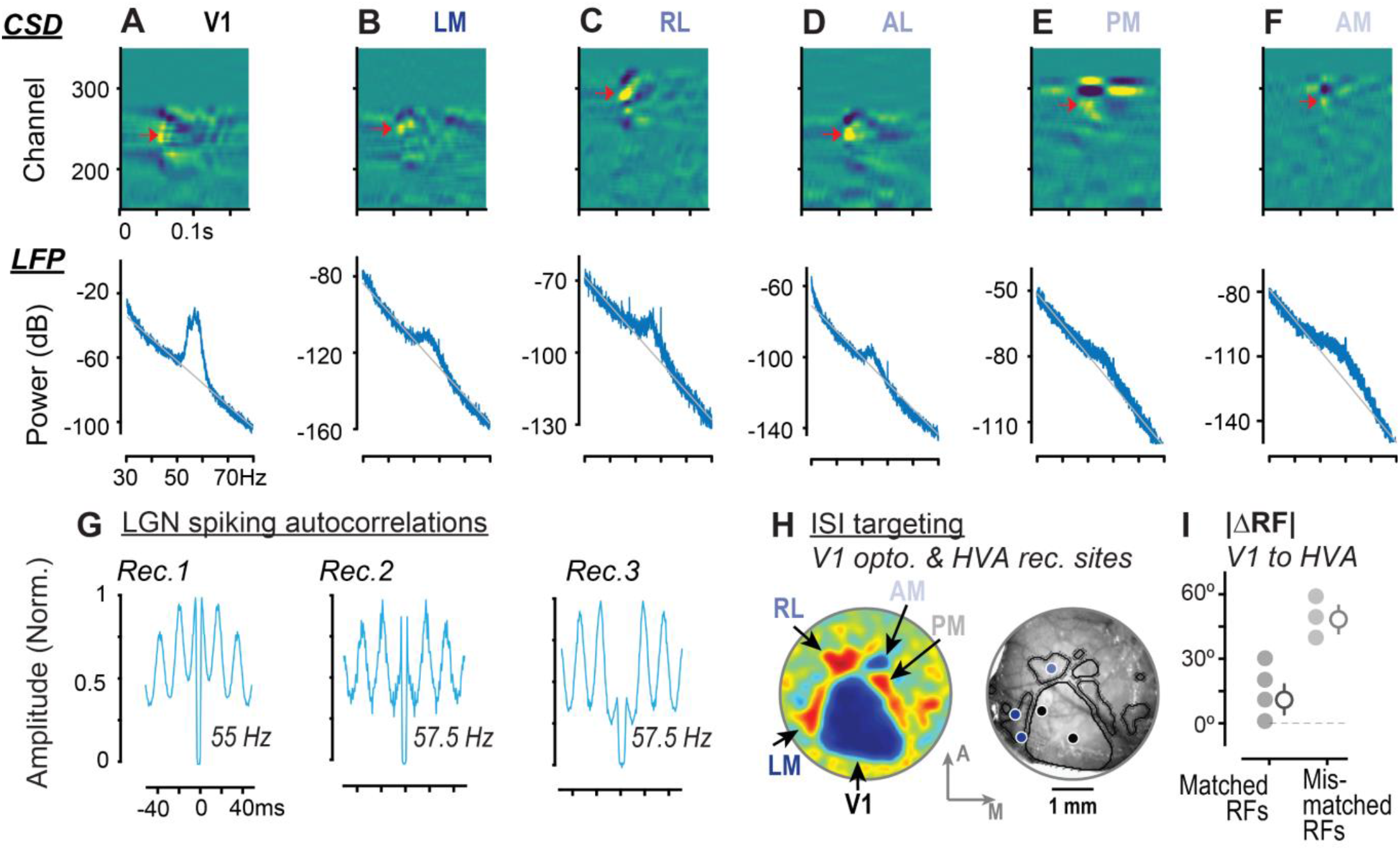
NBG LFP and CSD across cortical areas, LGN spiking and ISI targeting of V1 & HVAs. **A**. *Top*, Current source density (CSD) of local field potential (LFP) responses in an example V1 recording. CSD was computed with average LFP response to a full screen flash stimulus (onset at t = 0). Yellow shows current sink and indigo shows current source. Red arrow denotes channel with earliest sink, which corresponds to site with the earliest latency stimulus response (channel = 240; depth = 0.43 mm, latency = 38.4ms). *Bottom*, spectral power of spontaneous LFP activity at earliest sink site shows clear excess residual NBG power compared to 1/f exponential fit to spectrum (grey). Fit calculated by excluding the NBG range (50 – 70Hz) as in prior studies. Residual NBG power was considered in excess when mean power in NBG frequency range was > 1 SD above the 1/f fit. Recording session with 180 minutes of spontaneous activity; Another 10/14 V1 recordings showed excess spontaneous NBG residual LFP power compared to the exponential fit. **B**. Same as A, for lateromedial area (LM). *Top*, earliest sink channel = 252; depth = 0.31 mm; latency = 42.4ms. *Bottom*, example recording that showed excess NBG residual power versus exponential fit. 3/9 LM recordings showed excess spontaneous NBG residual power compared to the exponential fit. Note Y-axis scale vs A. **C**. Same as A, for rostrolateral area (RL). *Top*, earliest sink channel = 291; depth = 0.37 mm; latency = 42.4ms; *Bottom*, recording that showed excess NBG residual power versus exponential fit. 4/14 RL recordings showed excess spontaneous NBG residual LFP power compared to the exponential fit. **D**. Same as A, for anterolateral area (AL). *Top*, earliest sink channel = 241; depth = 0.38 mm; latency = 44.8ms. *Bottom*, recording that showed excess NBG residual LFP power versus exponential fit. 1/9 AL recordings showed excess spontaneous NBG residual power compared to the exponential fit. **E**. Same as A, for posteromedial area (PM). *Top*, earliest sink channel = 280; depth = 0.37 mm; latency = 50.4ms. *Bottom*, recording that showed excess NBG residual LFP power versus exponential fit. 2/10 PM recordings showed excess spontaneous NBG residual power compared to the exponential fit. **F**. Same as A, for anteromedial area (AM). *Top, e*arliest sink channel = 284; depth = 0.38 mm; latency = 56.8ms. *Bottom*, recording that showed excess NBG residual LFP power versus exponential fit. 2/9 AM recordings showed excess spontaneous NBG residual power compared to the exponential fit. **G**. Example auto-correlograms (ACGs) of identified NBG LGN neurons in 3 different recording sessions from Haider lab. Peak ACG frequency indicated for each neuron. Across 26 recordings in 7 mice, 196/805 (24%) of LGN neurons fulfilled NBG criteria (see Methods). **H**. Example intrinsic signal imaging (ISI) showing (*Left)* visual field sign map localizing V1 and multiple higher visual areas (HVAs), and (*Right)* craniotomies for recordings in LM or RL, and laser stimulation at two V1 sites. **I**. Experiments with matched azimuthal retinotopy at V1 stimulation and HVA recording sites (Δ10.2 ± 2.5°), versus mismatched sites (Δ47.4 ± 2.6°). Each data point indicates a single V1 stimulation and HVA recording pair (multiple recording sessions were conducted in each HVA with matched or mis-matched V1 stimulation).

**Figure S3.**
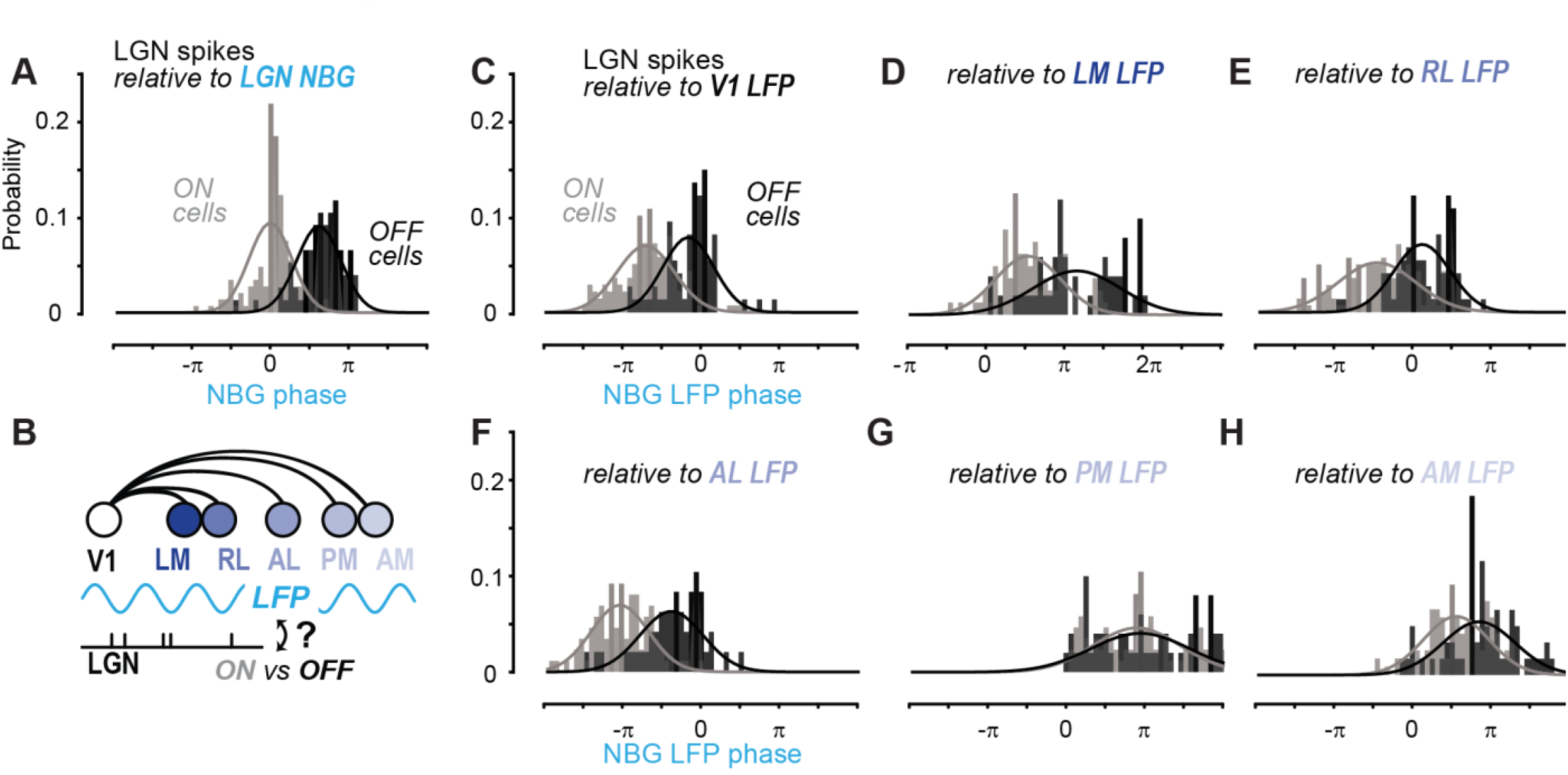
ON vs OFF LGN neuron spike phase relative to NBG LFP across visual cortical hierarchy. **A**. NBG phase of ON and OFF preference LGN neurons, relative to strongest NBG LGN ON neuron. Gaussian fits to ON/OFF cell phase histograms show clear preferred phase clustering and separation (ON cells: 0.016 ± 0.86 radians; OFF cells: 1.96 ± 0.89 radians; mean ± 1σ of Gaussian fit; n = 147 ON cells, 77 OFF cells, 14 recording sessions; see Methods for ON/OFF classification and phase calculations). **B**. Schematic of visual cortical anatomical hierarchy and relationship of LGN spiking in ON and OFF cells relative to cortical NBG LFP. **C**. Same as A, but LGN spikes referenced to simultaneously recorded V1 L4 LFP. NBG phase is identified with spike-LFP cycle histogram (as in Fig. 2E). ON cells: -2.13 ± 1.15 radians; OFF cells: -0.42 ± 1.02 radians; mean ± 1σ of Gaussian fit; n = 139 ON cells, 74 OFF cells, 13 recording sessions. **D**. Same as A but LGN spikes reference to simultaneously recorded LM earliest sink channel LFP. ON cells: 1.74 ± 1.31 radians; OFF cells: 3.70 ± 1.75 radians; mean ± 1σ of Gaussian fit; n = 79 ON cells, 50 OFF cells, 8 recording sessions. **E**. Same as A but LGN spikes reference to simultaneously recorded RL earliest sink channel LFP. ON cells: -1.44 ± 1.55 radians; OFF cells: 0.37 ± 1.13 radians; mean ± 1σ of Gaussian fit; n = 141 ON cells, 74 OFF cells, 13 recording sessions. **F**. Same as A but LGN spikes reference to simultaneously recorded AL earliest sink channel LFP. ON cells: -3.23 ± 1.15 radians; OFF cells: -1.17 ± 1.27 radians; mean ± 1σ of Gaussian fit; n = 87 ON cells, 48 OFF cells, 8 recording sessions. **G**. Same as A but LGN spikes reference to simultaneously recorded PM earliest sink channel LFP. ON cells: 2.86 ± 1.73 radians; OFF cells: 3.00 ± 1.97 radians; mean ± 1σ of Gaussian fit; n = 86 ON cells, 50 OFF cells, 9 recording sessions. **H**. Same as A but LGN spikes reference to simultaneously recorded AM earliest sink channel LFP. ON cells: 1.80 ± 1.31 radians; OFF cells: 2.71 ± 1.44 radians; mean ± 1σ of Gaussian fit; n = 117 ON cells, 59 OFF cells, 11 recording sessions.

**Figure S4.**
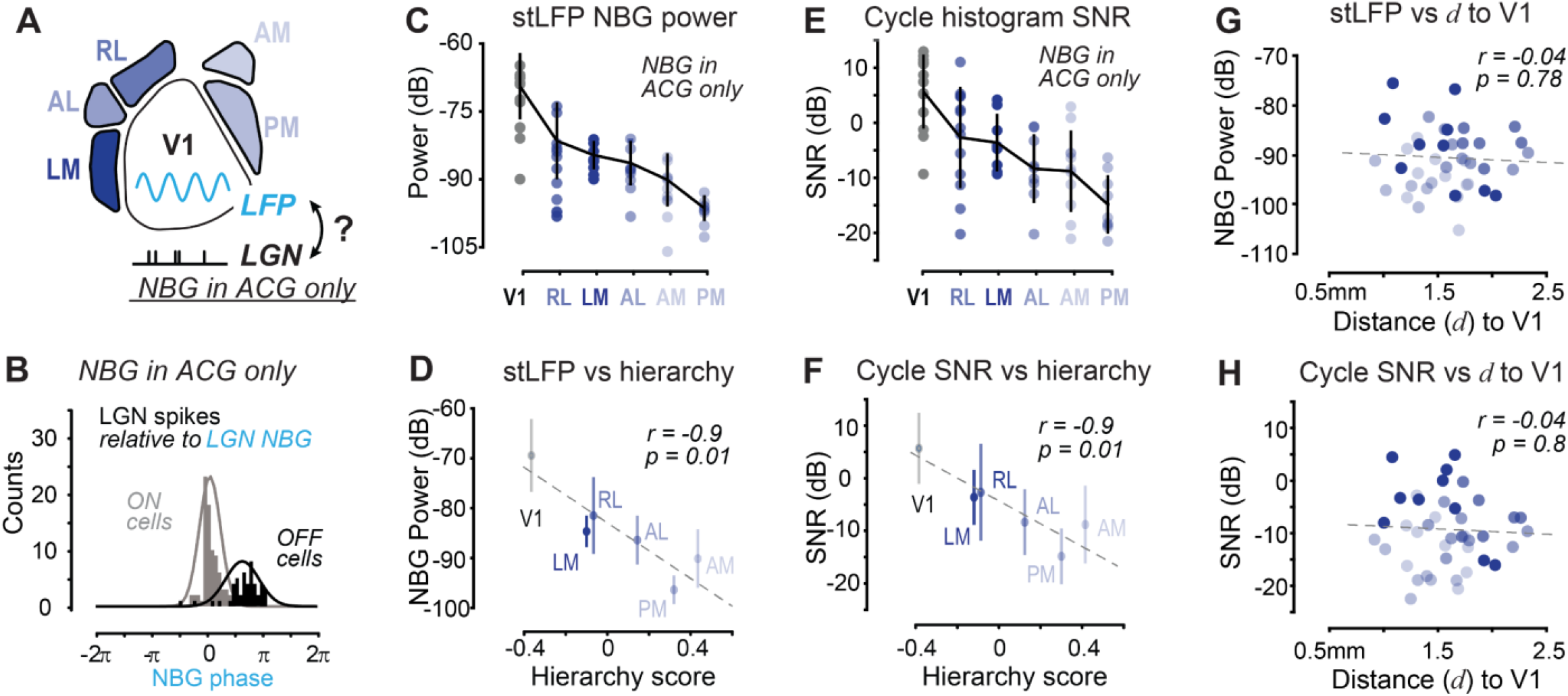
NBG neuron classification and volume conduction do not explain major findings. **A**. Schematic of LGN spiking relative to NBG phase of V1 L4 LFP, but only identifying NBG neurons in LGN based on spike auto-correlograms (ACG). **B**. NBG phase of ON and OFF preference LGN neurons, relative to strongest NBG LGN ON neuron. All NBG neurons classified solely with ACG criteria (n = 86 ON cells, 47 OFF cells, 14 recording sessions; see Methods), Gaussian fits to ON/OFF cell phase histograms again show clear preferred phase clustering and separation (cf. Fig. S3A). **C**. stLFP NBG power across V1 and HVAs using only LGN neurons with NBG in ACGs. (214 LGN NBG cells, 14 recording sessions). Errorbar shows mean ± SD, circles indicate individual sessions. Areas sorted from highest to lowest stLFP power. **D**. Same data as C, where stLFP NBG power of V1 and HVAs again shows strong and significant correlation with anatomical hierarchy values (Pearson rho = -0.90; *p = 0*.*0139*; see Methods for hierarchy scoring). **E**. Cycle histogram SNR of V1 and HVAs across all recording sessions, using only LGN neurons with NBG in ACGs. Same conventions as C. **F**. Same data as E, where NBG cycle histogram SNR shows strong and significant correlation with anatomical hierarchy. (Pearson rho = -0.90; *p = 0*.*0133*). **G**. stLFP NBG power of HVAs does not correlate with distance to V1. (Pearson rho = -0.04, *p = 0*.*78*; n = 41 recordings between V1 and HVA; 12 experiments, see Methods). In contrast, with the same data points, stLFP NBG power of HVAs shows strong and significant correlation with anatomical hierarchy. (Pearson rho = -0.42, *p* < *2e-3*) LGN NBG neurons identified with both ACG and CCG criteria, as in Fig. 3. See Methods for computation of distance of HVAs from V1. **H**. Similar to G, NBG cycle histogram SNR of HVAs does not correlate with distance to V1. (Pearson rho = -0.04, *p = 0*.*80*). In contrast, NBG cycle histogram SNR of HVAs shows strong and significant correlation with anatomical hierarchy. (Pearson rho = -0.44, *p* < *2e-3*)

**Table S1.**
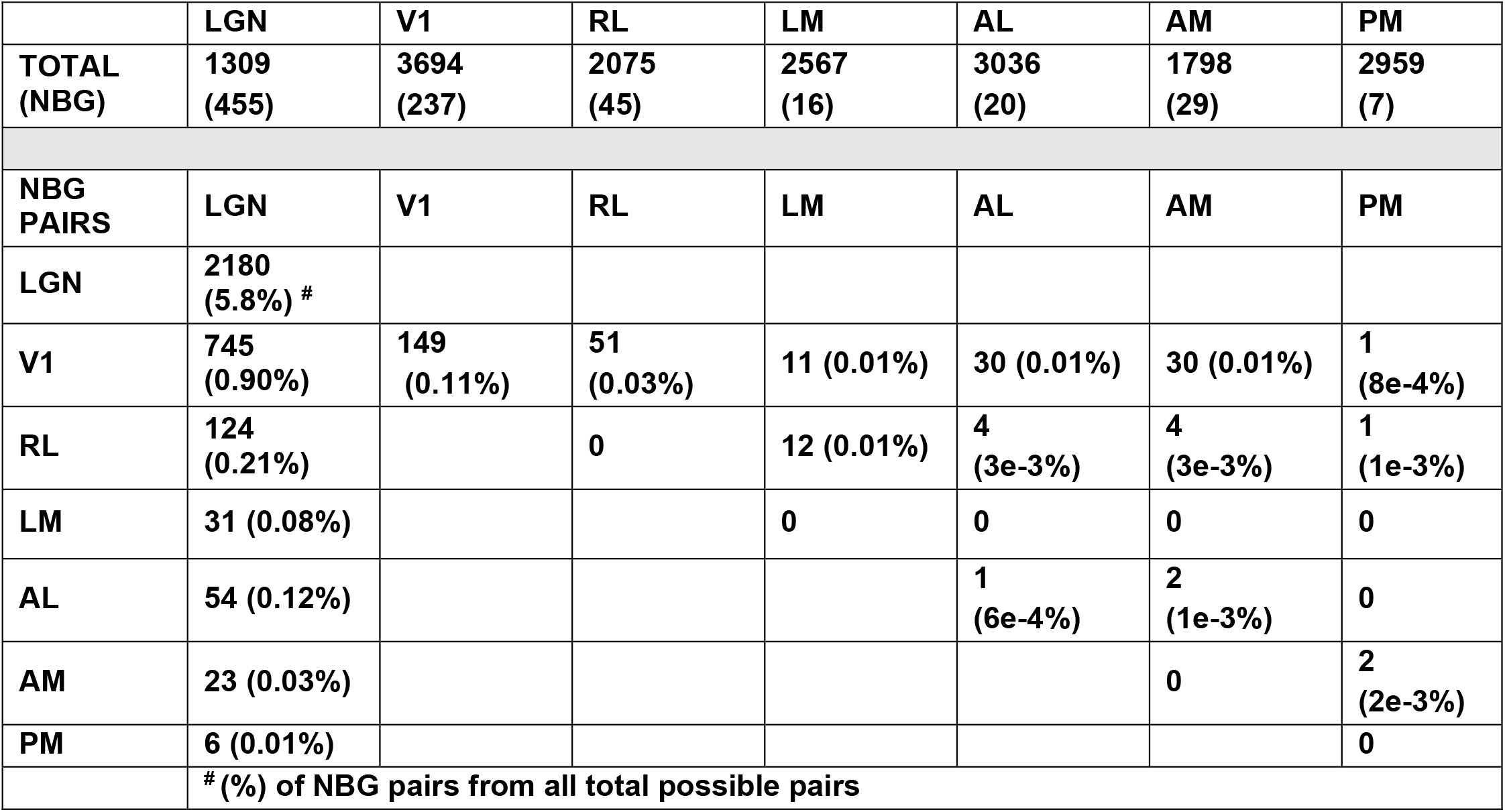
Number of NBG neurons (top) and NBG pairs exhibiting functional connectivity in CCGs between visual areas across all recordings in Allen Brain Observatory dataset. Top row indicates total number of neurons in each area (NBG neurons in parentheses). LGN: Lateral geniculate nucleus (32 recording sessions); V1: Primary visual cortex (56 recording sessions); LM: Lateromedial area (42 recording sessions); RL: Rostrolateral area (50 recording sessions); AL: Anterolateral area (44 recording sessions); AM: Anteromedial area (50 recording sessions); PM: Posteromedial area (36 recording sessions)

**Table S2.**
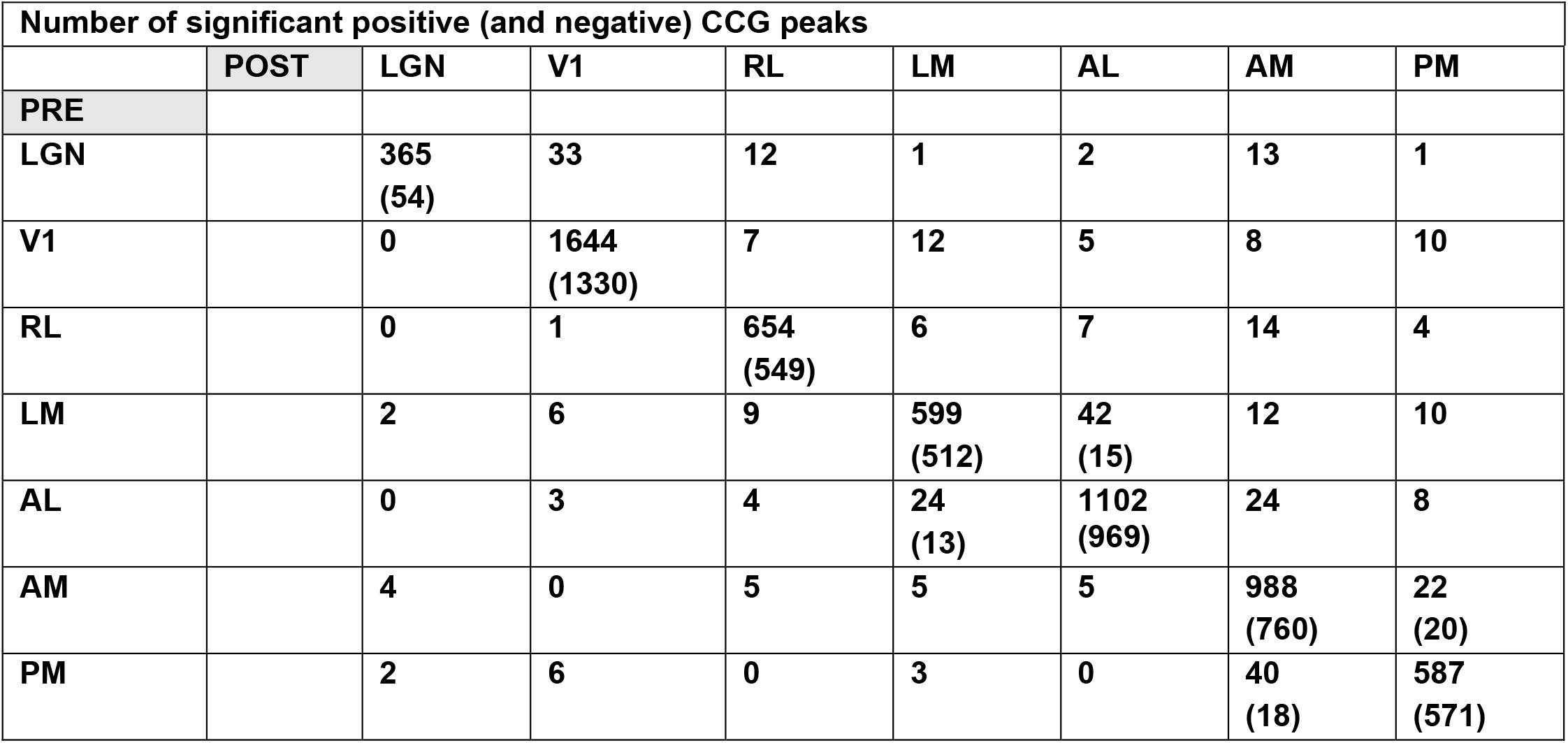
Number of pairs exhibiting positive (or negative) functional interactions within and across visual areas.

| Number of significant positive (and negative) CCG peaks |  |  |  |  |  |  |  |  |
| --- | --- | --- | --- | --- | --- | --- | --- | --- |
|  | POST | LGN | V1 | RL | LM | AL | AM | PM |
| PRE |  |  |  |  |  |  |  |  |
| LGN |  | 365<br>(54) | 33 | 12 | 1 | 2 | 13 | 1 |
| V1 |  | 0 | 1644<br>(1330) | 7 | 12 | 5 | 8 | 10 |
| RL |  | 0 | 1 | 654<br>(549) | 6 | 7 | 14 | 4 |
| LM |  | 2 | 6 | 9 | 599<br>(512) | 42<br>(15) | 12 | 10 |
| AL |  | 0 | 3 | 4 | 24<br>(13) | 1102<br>(969) | 24 | 8 |
| AM |  | 4 | 0 | 5 | 5 | 5 | 988<br>(760) | 22<br>(20) |
| PM |  | 2 | 6 | 0 | 3 | 0 | 40<br>(18) | 587<br>(571) |

**Table S3.** Cell-type specific probability of pairwise functional connectivity among high firing rate (High FR) & non-NBG (∼NBG) neurons (column 1), versus among NBG neurons (column 2). Cortical regular spiking (RS) and fast spiking (FS) neurons classified according to spike width (see Methods). Evidence for significantly enhanced probability of functional connectivity among NBG neurons (assessed with Binomial t-tests, see Methods). ^#^ LGN neurons with spike widths > 0.38 ms classified as “RS” (91% of all LGN neurons, 1187/1306; 91% of NBG LGN neurons, 415/455), assumed to be thalamocortical projection neurons.

| LGN to V1 functional connectivity |  |  |
| --- | --- | --- |
|  | P(FC \| High FR & ~NBG) | P(FC \| NBG) |
| LGN RS<sup>#</sup>-> V1 RS | 0%<br>(0/3513) | 0%<br>(0/2175) |
| LGN RS-> V1 FS | 0.05%<br>(1/2258) | 1.25% *<br>(26/2087)<br>*\*p < 1e-27* |
| LGN to HVAs functional connectivity |  |  |
|  | P(FC \| High FR & ~NBG) | P(FC \| NBG) |
| LGN RS-> HVA RS | 0.01%<br>(1/12626) | 0.70% *<br>(4/568)<br>*\*p<1.6e-7* |
| LGN RS-> HVA FS | 0.01%<br>(1/6977) | 1.25%<br>(13/1042)<br>*\*p < 1e-20* |
| V1 to V1 functional connectivity |  |  |
|  | P(FC \| High FR & ~NBG) | P(FC \| NBG) |
| V1 RS -> V1 RS | 0.47%<br>(46/9894) | 1.37% *<br>(14/1020)<br>*\*p<2e-4* |
| V1 RS -> V1 FS | 2.75%<br>(173/6300) | 5.92% *<br>(35/571)<br>*\*p<1.5e-5* |
| V1 FS -> V1 RS | 5.32%<br>(335/6300) | 7.28% *<br>(43/571)<br>*\*p<8e-3* |
| V1 FS-> V1 FS | 4.05%<br>(164/4054) | 7.36% *<br>(34/462)<br>*\*p< 3e-4* |

